# Lack of detectable neoantigen depletion in treatment-naïve cancers

**DOI:** 10.1101/2023.06.21.544805

**Authors:** Maarten Slagter, Lorenzo F. Fanchi, Marit M. van Buuren, Arno Velds, Gergana Bounova, Lodewyk F.A. Wessels, Ton N. Schumacher

## Abstract

While neoantigen depletion, a form of immunoediting due to Darwinian pressure exerted by the T cell based immune system during tumor evolution, has been clearly described in murine models, its prevalence in treatment-naive, developing human tumors remains controversial. We developed two novel methodologies to test for depletion of predicted neoantigens in patient cohorts, which both compare patients in terms of their expected number of neoantigens per mutational event. Application of these strategies to TCGA patient cohorts showed that neither basic nor more extensive versions of the methodologies, controlling for confounding factors such as genomic loss of the HLA locus, provided statistically significant evidence for neoantigen depletion. In the subset of analyses that did show a trend towards neoantigen depletion, statistical significance was not reached and depletion was not consistently observed across HLA alleles. Our results challenge the notion that neoantigen depletion is detectable in cohorts of unmatched patient samples using HLA binding prediction-based methodology.

## Introduction

Immune evasion, the avoidance of immune detection and eradication, is a hallmark of cancer^1, 2^ and can occur through various mechanisms, including deletion of components of the antigen presentation machinery, insensitivity to pro-apoptotic or growth-arresting molecules, such as granzymes and IFNg^3^, or expression of T-cell checkpoint ligands such as PD-L1. In addition, the Darwinian selective pressure exerted by T cells has been proposed to lead to outgrowth of tumor cells that lack T cell-recognized (neo–)antigens. In line with this, loss of a neoantigen that was recognized by T cells has been observed in a murine sarcoma model^4^. In addition, loss and reduced expression of mutant genes encoding T-cell recognized neoantigens has been observed in two patient case studies^5^, and the reduction in tumor mutational burden observed in clinical responders to PD-1 blockade has been proposed to lower the number of T cell-recognized neoantigens^6^. While collectively these data form relatively strong evidence that neoantigens can be lost upon (therapeutically enhanced) T-cell pressure, it is less clear whether such genomic neoantigen depletion of neoantigens widely occurs in treatment-naïve tumors, and – importantly – would be detectable at the genomic level. An argument against the idea that neoantigen depletion could be readily detected at the genomic level is formed by the observation that only a minor fraction of predicted neoantigens appears to naturally induce T-cell responses in patients^7–10^. Importantly, only this small subset of predicted neoantigens can be expected to be under Darwinian pressure, and this will affect the sensitivity of any methodology that examines the occurrence of predicted neoantigens regardless of the fact whether or not T cell reactivity was present against these predicted antigens.

Several prior studies have assessed neoantigen depletion in large sets of cancer genomes with unmatched samples (i.e., with a single tumor sample per patient) provided by the Cancer Genome Atlas (TCGA). A first of these studies provided evidence for the selective loss of mutations predicted to encode MHC-class I neoantigens for colorectal and clear cell kidney cancer^11^, but at the same time observed a counter-intuitive neoantigen enrichment in EBV^-^ stomach adenocarcinoma. Likewise, application of a model of peptide immunogenicity found recurrent mutations to appear more readily in TCGA patients that are less capable of presenting the resulting new peptide sequences by HLA^12^. In line with these results, an earlier TCGA pan-cancer analysis found recurrent oncogenic mutations to be relatively poorly HLA-presentable by the patients that carried them^13^, a result that has however since been shown to be fully driven by confounding factors^14, 15^. The same group also reported elevated neoantigen levels in tumors that harbor mutations in the antigen presentation pathway, but this result was not corrected for, potentially confounding, background mutation rates^16^. Using a mathematical model of tumor evolution, Lakatos et al. predicted the variant allele frequency (VAF)-spectrum of a tumor’s somatic mutations under various degrees of immune pressure and found TCGA tumors to appear similar to simulations under immune pressure^17^. In contrast to the aforementioned studies, a comparison of the ratio of non-synonymous to synonymous mutation count (‘dN/dS’) between areas of the human genome that do or do not encode predicted HLA presented peptides found no evidence of negative selection when correcting for sequence content between these two classes of genomic regions^18^. Notably, this sequence content was shown to affect the ratio with which mutagenic processes yield either non-synonymous or synonymous mutations and so confounded dN/dS estimates^18^. A more recent study using dN/dS methodology did report depletion, especially of clonal mutations, in highly immune infiltrated tumor types and in patients that did not show other means of immune evasion, but did however not correct for sequence content^19^. This latter study also showed substantial degrees of nonsensical neoantigen enrichment, especially in lowly immune infiltrated tumor types.

Given the conflicting results in these prior analyses, there is a need for novel methodology to assess neoantigen depletion in treatment-naïve tumors. Here, we present two new, interrelated, methods for the detection of average neoantigen depletion in patient cohorts. Using these methods, we do not observe substantial evidence for neoantigen depletion in TCGA tumors, despite incorporation of many potential confounding co-variates into our modeling to increase sensitivity. We emphasize that the lack of signal that we describe does not rule out the occurrence of neoantigen depletion for a minority of studied patients and/or a minority of (T cell-recognized) neoantigens. However, our observations do challenge the notion that neoantigen depletion signals based on HLA affinity predictions are detectable in large-scale unmatched cancer sequencing data sets.

## Materials & Methods

### Datasets

#### HIV peptide data for pipeline prediction validation

We compiled a list of 81 HLA-A*02:01-restricted HIV T-cell epitopes by querying the Los Alamos National Security HIV Database (https://hiv.lanl.gov, December 2010) and PubMed. The obtained peptides were filtered according to the following criteria to create a list of ‘high confidence’ HLA-A*02:01-restricted and T-cell recognized epitopes: (1) epitope presentation was shown to be restricted to the HLA-A*02:01 allele; (2) T-cell reactivity against the epitope was reported in at least 3 patients/studies, and (3) evidence of endogenous processing of the epitope had been obtained (i.e., T-cell responses observed in vaccination or peptide loading studies were excluded). These criteria were met by 32 of 81 acquired peptides.

To assess all candidate HIV-1 epitopes, we acquired the assembled sequencing data of an HIV-1 isolate from the NCBI database (isolate 671-00T36; NCBI accession number AY423387^1^) and considered it a reference HIV-1 genome. Partitioning this reference genome in all nonameric peptides, 3,094 candidate peptides were generated. Out of the 32 ‘high confidence’ HIV epitopes discussed above, 17 were not perfectly mappable to the reference genome (1-2 amino acid differences at most). To correct for this and allow cross-matching, the reference genome was adjusted to exactly match the mismatching peptide sequences (adjusted reference in Table S2). One out 17 epitopes remained completely unmappable to the reference genome and was thus excluded from further analysis (Table S2).

#### IEDB peptide data for pipeline validation

Peptides selected for T cell recognition proven usen any methodology were downloaded from the Immune Epitope Database (www.iedb.org, 2018-12-10) from the ‘Assays’ section. Assays were filtered for: i) having four-digit HLA-typing (e.g., ‘HLA*B-27:05’), ii) having the targeted peptide be available and 9 amino acids of length in the (Antigen Description field) iii) having an entry for the Organism species name. Peptides were considered T-cell targetable if at least 2 tested subjects responded, combined over all the assays investigating a particular peptide. If this field was not available for any of the assays for a particular peptide, the peptide was considered T-cell targeted if at least one assay gave a positive result (i.e., response labelled as Positive, Positive-High, Positive-Intermediate or Positive-Low). Notice that these criteria are substantially less stringent than the ones employed for the focused HIV-set. This analysis was restricted to HLA-A*02:01 peptides as this allele was by far the best represented in the IEDB. Next, redundancy in the acquired T-cell targeted peptide sets per pathogen was removed by iterative sequence alignments of all unordered peptides against all remaining peptides using the function `pairwiseAlignment` from the R Bioconductor package Biostrings (version 3.8). Matching scores above zero were interpreted as sequence similarity, resulting in removal of the second peptide of the pair from the peptide set. This way, the T-cell targeted peptides in the resulting list were all dissimilar from each other. Next, in order to predict proteasomal processing efficiency of candidate peptides, the amino acid context of the peptides in their source protein was required, but this information was not included for all peptides in the IEDB. To obtain this information, peptides were mapped to the reference proteomes of the viruses they were annotated to originate from using `phmmer` (version 3.2.1) with the –`max` and – `domtblout` flags, prioritizing matching reference sequences by their alignment length (the number of matching amino acids between query and target sequence, longer alignments preferred), the alignment discrepancy (when available, the difference between the annotated C-terminus and the inferred one, smaller is preferable), the source of the reference sequence (the manually curated SwissProt prioritized over the more exhaustive TrEMBL), the alignment’s *e*-value and the query name, in that order. Query peptides with more than 3 mismatches between the source and query sequences were excluded from further analysis. The following viruses reference proteomes were used: Human gammaherpesvirus 4 (EBV, UP000007639), Human Immunodeficiency Virus 1 (HIV-1, AUP000002241), Alphapapillomavirus 9 (AUP000009104), Vaccinia virus (AUP000000344), Influenza-A virus (AUP000131152), Hepatitis-B virus (AUP000008591), Hepacivirus-C (AUP000000518), Human alphaherpesvirus 1 (AUP000106517), Dengue virus (AUP000002500), Human betaherpesvirus-5 (AUP000000938). Transcriptome references were downloaded from the UniProt database by querying for the virus name and downloading all (possibly redundant) proteins (The UniProt, 2017). For non-perfectly mapping peptides, the most highly rated reference sequence was then modified to reflect the query sequence (i.e., peptide), such that the most representative processing score for the peptide could be computed.

#### TCGA data

The patient sample cohort, consisting of all tumor samples for which both DNA, RNA sequencing data was available, spans 5,585 patients from 30 tumor types (Table S1). TCGA data acquisition from the Broad Institute’s Firehose and integration and preprocessing of data sources listed below was automated in R using functionality that is combined in the R package `firehosedownload` (github.com/slagtermaarten/firehosedownload).

TCGA somatic variant calls in MAF-format and RNA sequencing data were downloaded and harmonized from the 2015-08-21 release of the Broad TCGA genome data analysis center standard runs (http://gdac.broadinstitute.org/runs/stddata). As mutation data for the ESCA (oesophagus carcinoma) project is not part of this release, mutations for the ESCA project were obtained from the repository of mutations files curated by Cyriac Kandoth (http://www.synapse.org/#!Synapse:syn1695396.13). TCGA RNA sequencing data were downloaded from the Broad TCGA genome data analysis center 2015-11-01 release of the standard runs (http://gdac.broadinstitute.org/runs/stddata).

For projects where data from multiple sequencing platforms were available, Illumina HiSeq data was preferentially used. Raw read counts in RNAseq data were subsequently transformed to the transcripts per million (TPM) RNA abundance measure, using custom R functionality included in `firehosedownload` and using Ensembl75 (release of February 2014) gene length information.

#### TCGA HLA typing data

HLA typing based on DNA sequencing using OptiType^3^ was downloaded from the TCIA resource (www.tcia.at) provided by the Trajanoski laboratory.

### Genome– and patient-level annotation

#### MMR status of tumor samples

The R package MSIseq^4^ was used to infer microsatellite instability status of all TCGA tumor samples in our cohort. Mutation annotation format files were obtained from TCGA as described above. Sequencing target region sizes were calculated for each sample from target enrichment design files used in the various TCGA projects (Supplementary Table S3). These data were subsequently used as input for the MSIseq classifier.

#### Annotation of antigen presentation capability and T cell sensitivity of tumor samples

To identify tumor samples that may be resistant to T cell attack, we analyzed samples for occurrence of non-synonymous mutations in any of the 515 non-HLA genes identified to potentially induce resistance to CD8^+^ T cell mediated killing^5^, or for occurrence of non-synonymous mutations in any HLA class I allele^6^.

#### PAM50 subtyping of breast cancer samples

We downloaded Level 3 RNA-seq data for the BRCA cohort from the TCGA Data Portal on 2015-06-25 and analyzed the expectation maximization normalized counts. Samples were PAM50-subtyped independently^7^ using the implementation in the genefu R package^8^ with the robust scaling option enabled.

#### Allele-specific HLA loss

We adapted the tool LOHHLA^9^, which allows for allele-specific detection of genomic aberration of HLA alleles, to make it more amenable to large scale application (code available on GitHub: https://github.com/slagtermaarten/LOHHLA). Most importantly, we included support for the reference genome GRCh38, made it compatible with single-end sequencing data and expanded it such that it can handle input bam files that are restricted to the HLA region of interest, rather than whole exome or whole genome bam files. The original version of LOHHLA compares whole-genome or whole-exome coverage mapped reads (bam formatted) between a tumor and matched normal sample, comparing read coverage normalized to the total number of mapped reads between the two samples for each HLA allele separately. To circumvent having to download the full bam files for thousands of patients just to obtain the total number of mapped reads per bam file, we inferred the total number of mapped reads from the file sizes of the complete bam files (accessible from the NIH GDC API). For this an ordinary linear regression model was used, which was fitted on file sizes in bytes and the number of mapped reads (as read out using `samtools` `flagstat`, 5^th^ row of output) of 114 normal and tumor sample bam files for 60 randomly sampled patients (Table S5) for which we did download the entire bam file using `gdc-client`, deriving the expression: `total` `mapped` `reads` `=` 88370554 + 7461 [reads/MB] * file size [MB] (Figure S6A). This allowed us to use the TCGA GDC bam slicing API (https://api.gdc.cancer.gov/slicing/view/) to specifically download the HLA regions of chromosome 6 (29941260-29945884, 31353872-31357187 and 31268749-31272092 in GRCh38 coordinates) and use those as input for LOHHLA. Note that the full TCGA bam files that were obtained largely (∼99.9%) consisted of mapped reads, obviating the need to correct for the presence of unmapped reads. In order to ensure that read pairs for which one of the read mates lied outside the annotated HLA genomic range were included in the analysis, the bam slicing download window was extended by 10^7^ bp on both the 3’ and 5’ sides to 28941260-32357187. We fed the LOHHLA analyses purity and ploidy estimates obtained from ASCAT as described in the section ‘Variant cellularity’ below. The minCoverage argument to LOHHLA determines the minimally required amount of coverage in the normal sample in order for an SNP to be considered eligible to contribute to the copy number estimate of the gene it’s positioned in. It was set to 0, after having tested the stability of the copy number estimates by titrating the minCoverage filter between 0 and a patient specific upper boundary computed as the median coverage of sites in the matched normal sample. While titrating the minCoverage threshold, we recorded the inferred copy number for each allele (i.e., `HLA_type{1,2}copyNum_withBAF,` which is computed as the median of the copy number estimates for individual loci pertaining to an allele, Figure S6B). Next, we computed the coefficient of variation (CoV, standard deviation/mean) over the copy number estimates to obtain donor, gene, and allele-specific estimates of the effect of the minCoverage threshold on the copy number estimates (Figure S6C). Alleles that showed a CoV < .25 were labelled as robustly estimable (90% of alleles, Figure S6D), resulting in 63% of evaluated patients to have robust estimates for all of their HLA-A, –B, and –C alleles (Figure S6E). Immunoediting analyses with the ‘strict LOHHLA’ attribute included only these patients; analyses with the ’lenient LOHHLA’ attribute also included patients for which not all alleles were robustly estimable. An overview of allele-specific copy number estimates for HLA-A, –B, and –C are displayed in Figure S6F. Alleles were deemed lost if the upper boundary of the copy number estimate’s 95% confidence interval was below 0 and the estimate’s minCoverage CoV was below .25.

#### Cytolytic score

Cytolytic score^10^, a transcriptomic proxy for T and NK-cell activity, was computed as the geometric mean of TPM-normalized expression estimates of the *PRF1* and *GZMA* genes: exp(0.5 ln(PRF1 + 0.01) + 0.5 ln(GZMA + 0.01))

### Gene-level annotation

#### Gene essentiality

To infer the essentiality of genes to cell survival, we integrated the work of the Sabatini^11^ and Brummelkamp^12^ laboratories. Wang et al.^11^ screened the Jiyoye and Raji cell lines (Burkitt Lymphoma) and the K562 and KBM7 (CML) cell lines using CRISPR technology, and the KBM7 cell line using GeneTrap technology. Blomen et al.^12^ screened the KBM7 cell line and its derivative, the HAP1 cell line, using GeneTrap technology. Blomen et al. provided binary class labels indicating essentiality for genes, whereas Wang et al. only provided raw read count data, presenting significance testing for only a subset of performed experiments.

In the Wang et al. CRISPR KBM7 data analysis, essential genes were considered to have a CRISPR-score (average log_2_ fold change in the abundance of sgRNAs targeting the gene) lower than –0.1 and an adjusted *p*-value below 0.05. Neither general nor cell line-specific criteria were included for the other three cell lines that Wang et al. screened using CRISPR technology. We elected to apply the same criteria to the other three cell lines screened with CRISPR technology by Wang et al. Similarly, no cut-off was proposed for the Wang et al. GeneTrap data set, but rather a correlation between the GeneTrap and CRISPR data was reported. Here, we used a minimum required amount of anti-sense inserts of 65 and set the required GeneTrap score to be lower than .45, in order to maximize the similarity in set cardinalities between the Sabatini-KBM7-GeneTrap and the Sabatini CRISPR/KBM7 derived gene set (1,875 and 1,878 genes, respectively). Having obtained seven partially overlapping lists of essential genes, any gene appearing at least once in any of the seven experiments was deemed essential, 10.6% of selected genes appeared in all lists. The list of genes is included in Table S4.

### Peptide-level annotation

#### Similarity to self-repertoire

To determine whether predicted neoantigens were likely to be dissimilar enough from self to be recognized by the endogenous T-cell repertoire, we implemented a self-similarity classifier based on previously identified determinants of T cell similarity^13^. We compare candidate epitopes arising from somatic mutations to peptides from the human proteome predicted to be presented by the relevant HLA allele (e.g., HLA-A*02:01), restricting this comparison to the amino acids spanning positions 3 to 8 of nonameric peptides, as these are considered to be most important for T-cell recognition of peptide-HLA complexes^13–15^. Epitopes were deemed ‘dissimilar-from-self’ when one or more of the following criteria are met: (1) amino acid position 5 is mutated, (2) T-cell exposed region contains 3 or more mutations, (3) two mutations are clustered to one side of position 5 (i.e., in positions 2-3-4 or positions 6-7-8), and (4) a single amino acid substitution leads to large physicochemical changes on position 2, 3, 4, 6 or 7. The latter substitutions were defined as amino acid changes with an absolute covariance of < 0.05 in the PMBEC amino acid similarity matrix^16^. We implemented this algorithm in Rcpp (C++) and distributed it as part of the quickMHC R package (www.github.com/slagtermaarten/quickMHC).

### Somatic variant annotation

#### Variant effect prediction

To determine the effects of the various classes of mutations found in tumor samples, we developed a Perl tool named VarContext. Canonical cDNA transcripts were obtained for genes containing mutations (single nucleotide variants and/or indels) by querying the Ensembl database (release 75, GRCh37). We applied each mutation affecting a particular gene and annotated its effect (silent, missense, insertion, deletion, frameshift, stop loss, stop gained), yielding the tumor transcript sequence. Transcripts which lost their stop codon as a consequence of the applied mutations were extended until the next in-frame stop codon was encountered. In contrast, transcripts gaining a premature termination codon (PTC) were analyzed for their potential of triggering nonsense-mediated decay (NMD; detailed description in ‘Transcript-level annotation’). Finally, the canonical and modified cDNA sequences were translated into amino acid sequence, resulting in the reference and tumor transcripts.

#### Variant oncogenicity and essentiality

The oncogenic potential of mutations was predicted using IntOGen mutations^17^ version ‘IntOGen Mutations Analysis 2.4.1-maintenance’ (https://www.intogen.org/analysis), specifically for all affected transcripts and stratified by sequencing project. TCGA maf files were converted to the required TSV-based input file format using a custom script and were analyzed in December 2015. Variants were further annotated with a driver gene score indicating the likelihood of being a driver gene based on the gene of their location^18^. To infer essentiality to cell survival of individual genes, we integrated the work of two laboratories^11, 12^ into an overall essentiality score, with which somatic mutations were subsequently annotated (see ‘Gene essentiality’). Additionally, annotation of oncogenicity was performed by comparing gene, amino acid change and amino acid change position (determined using VarContext) of somatic mutations to the list of 1,018 recurrent oncogenic mutations compiled by Marty et al.^19^. The latter list was used for the ‘Marty’ variant selection setting.

#### Variant cellularity

Clonal antigens that are presented by all tumor cells in a lesion can reasonably be expected to have a larger contribution to tumor regression than subclonal antigens, and the observed inverse relationship between tumor heterogeneity and immunotherapy outcome provides indirect support for the superior value of clonal antigens as T-cell targets ^20^.

We estimated the cellularity of individual mutations from DNA sequencing read count information for a subset of 3,660 tumors in 18 tumor types, selected based on availability of the required data types. As a starting point for inferring the fraction of tumor cells that carry a mutation (the variant’s cellularity), we use the variant’s observed allelic frequency (VAF), the fraction of reads overlapping with the variant locus that carries the variant. The observed VAF of a somatic variant does not relate to its cellularity in a straightforward way as it is a compound measure of several factors: the proportion of contaminating normal cells, the number of allelic copies of the variant in each cell and its cellularity, plus uncharacterized sources of technical noise^21^. We employed two methods of inferring variant cellularity. The first is based on published maximum likelihood-based approaches^20, 22^, the second is a Bayesian hierarchical clustering of variants^21^ for potentially more robust and accurate cellularity estimates.

First, variants were annotated with the absolute copy number status of the genomic segments they are located in. Absolute copy number status was derived from ASCAT analyses of Affymetrix SNP6 profiles^23^ and were obtained from the COSMIC resource (https://cancer.sanger.ac.uk/cosmic/download) on 2016-04-24. Absence of coverage in the ASCAT file was assumed to imply absence of local copy number aberrations as only small parts of the genome are covered in the SNP6 output. Additionally, tumor purity estimates, representing the percentage of tumor cells in the sample, were obtained from these ASCAT analyses. The intersection of patients eligible for neoantigen prediction (patients for which DNA and RNA sequencing data was available) and those for which read count information and a COSMIC ASCAT analysis was available (required for cellularity estimates) consisted of 3689 TCGA patients, distributed over 13 sequencing projects (see Supplemental Table 1 for the exact list of included samples).

#### Maximum likelihood-approach for cellularity estimation

Following Landau et al.^22^, we note that the observed number of reads consistent with the called mutation, *N_a_*, is binomially distributed: *P*(*N_a_*) ∼ *Bin*(*N*, *AF*_e_(*c*)), where *N* denotes the total number of reads covering the genomic locus of the variant and *AF_e_* denotes the expected allele fraction of the variant under a particular fraction of cells carrying the mutation *c* (for cellularity), on which *AF_e_* depends as follows:

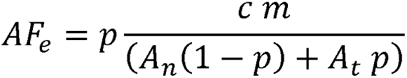

In which *p* denotes the tumor purity of the sample, i.e., the fraction of cells that are cancer cells, *A_n_* and *A_t_* denote the average amount of alleles in the normal and tumor populations, respectively, and *m* denotes the variant’s multiplicity, i.e., the number of tumor alleles that carry the variant – an integral number smaller than or equal to *A_t_* assumed to be equal across all tumor clones. The listed expression for *AF_e_* can be understood as the fraction of the number of tumor cell alleles carrying the mutant allele and the total number of alleles, from both tumor and normal cells, at the somatic variant’s genomic locus.

Two of the quantities on which *AF_e_* depends are not directly observed: *c* and *m*. Thus, neither *c* nor *m* are unambiguously identifiable without knowledge or assumptions about the other. In the case where the *VAF* equals the fraction of alleles derived from tumor cells 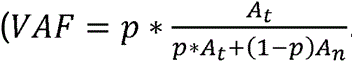, only one scenario is plausible: *c* = 1 and loss-of-heterozygosity (LOH) must have occurred at the variant loci in the cell giving rise to all sequenced cells (*m*= *A_t_*). Ambiguity however arises for mutations located in genomic regions of copy number aberrations and for which the *VAF* does not unequivocally indicate m to equal *CN_t_* (i.e., *VAF* < 1). This ambiguity is caused by the fact that a subclonal mutation (low cellularity) with high multiplicity could result in similar VAF-values as a clonal variant (high cellularity) with low multiplicity. Following the examples of Landau et al. and McGranahan et al. ^24^, implicit in the case of McGranahan et al., we assume the multiplicity of the variant to be unity when estimating cellularity, thereby running the risk of overestimating the *c* of somatic variants for which the multiplicity potentially exceeds one, i.e., variants located in an amplified segment. For some variants, it is certain that the multiplicity must have been greater than unity as *m*= 1 results in *VAF*)*AF_e_* for *c*= 1. For these, we iteratively increase *m* by 1 until *VAF* falls in the range of expected allelic frequencies (*AF_e_*), stopping before *m* exceeds the major allele count. Having set the multiplicity *m* to a minimal value consistent with the observed data, we can proceed to compute the most likely cancer cell fraction *c*. Assuming a discrete uniform prior on *c* in the range [0,1], discretized in intervals of 1/1,000, we compute the likelihood of each *c* under the binomial model *P*(*c*) and subsequently normalize the likelihoods by dividing them by their sum – the constant of proportionality of *P*(*c*) – to obtain the posterior probability mass function of *P*(*c*)^22^. The mode of this distribution, i.e., the maximum likelihood estimate is taken as the somatic variant’s cellularity, with the two boundaries centered on the mode and encompassing 95% of the distribution as the 95% credibility interval. Following McGranahan et al., we labelled mutations as clonal if their inferred *c* was greater than or equal to 0.95 and the upper boundary of the *c* 95% credibility interval was greater than or equal to .99.

#### PyClone for cellularity estimation

In addition to the ML-based approach, we ran PyClone^21^ for all samples with read count information, ASCAT purity and absolute copy number estimates available and a median total read count on called variants exceeding 100 reads (simulations succeeded for 780 tumors, see Supplemental Table 1 for the list of included patients). We ran the binomial model for 10^5^ Markov Chain Monte Carlo iterations, using a Markov Chain burn-in of a 1,000 iterations and Markov Chain thinning of 5.

#### Correspondence between the two methods

We hypothesized that the largest structural difference between the two methods lies in the inference of allelic frequency, which is only of relevance for mutations located in genomic regions with copy number aberrations. In support of this, the two methods yield highly similar and correlated clonality estimates for those mutations that do not require an inference of allele multiplicity (Figures S2A and S2B). As this class forms the large majority of mutations (98.75% of somatic variants), further analyses were performed using the maximum likelihood-based approach, because of its more clearly defined dependency on sequencing depth. To provide a validation of the obtained cellularity estimates, we compared cellularity estimates between the most highly recurrent mutations within a given tumor type and the aggregate of non-recurrent mutations. Consistent with expectations, for the majority of tumor types (15 out of 16) the most highly recurrent mutation was predicted to be significantly more clonal than the remainder of the mutations observed in that tumor type (Figure S2C). As we deemed the correspondence between the two methods satisfactory, further analyses were based on the ML-based method for its relative simplicity, computational efficiency and relatively low requirements on read coverage.

#### Variant-specific expression

To estimate the expression of individual somatic variants, we required the RNASeq mapped reads from the TCGA GDC which are mapped to GRCh38.d1.vd1. We used the bam slicing tool in order to prevent having to download entire .bam files. We first converted the variant loci from the hg19 to GRCh38 coordinate system, using the `rtracklayer` and `Granges` functions out of the `rtracklayer` (version 1.42.2) and `GenomicRanges` (version 1.34.0) R Bioconductor packages in combination with a liftover file obtained from UCSC (ftp://hgdownload.cse.ucsc.edu/goldenPath/hg38/liftOver/hg19ToHg38.over.chain.gz). These transformed coordinates were subsequently used to query the TCGA GDC bam slicing tool, using a random UUID in case multiple UUIDs were listed for RNASeq of tumor samples from the patient. Next, the number of reads consistent with the variant and reference alleles was tallied by running `samtools` `mpileup` (version 1.9) and custom R functionality.

### Neoantigen prediction

Tumor transcripts were reconstructed from SNVs and indels in order to obtain a set of candidate tumor-specific neoantigens. Candidate peptides whose genomic sequences were affected by multiple mutations were modified to reflect the consequences of all variants (1.75% of all candidate peptides). SNVs and frameshifting insertion-deletions can introduce premature termination codons (PTCs), rendering the encoded transcripts prone to degradation by the nonsense-mediated decay (NMD) machinery. To account for this, we implemented an NMD-calling routine based on previously inferred characteristics of NMD-targeted transcripts^25^. As expected, PTC-inducing variants that were classified as invoking NMD had significantly lower variant allele fractions (VAF) in RNA sequencing data than in DNA sequencing data, indicating degradation of PTC-bearing transcripts (Figure S1G). Hence, NMD-predicted transcripts were removed from further analysis.

Reference and tumor transcripts were used as inputs for Neolution, the in-house neoantigen prediction pipeline that annotates tumor-specific transcript derived peptides with four scores representing the various stages of antigen presentation, classifying those passing all four filters as MHC-binding peptides likely to yield an immune response. We assess the following steps: (i) RNA expression, (ii) proteasomal processing and transport into the endoplasmatic reticulum, (iii) MHC binding, and (iv) dissimilarity from self-antigens. First, we determined whether genes encoding candidate neoantigens are expressed, excluding – unless indicated otherwise – all peptides for which the associated gene had an expression level surpassing a constant threshold (default: 0). Alternatively, we applied a threshold to library size-normalized read counts at the variant level as described in ‘Variant expression’. Second, we used netChop^26^ to predict the likelihood of successful peptide processing by the proteasome and TAP transport (NetChop score ≥ .5). Third, we used netMHCpan3.0^27^ to predict HLA class I binding affinity, by default employing a percentile rank threshold of 1.9 (which corresponds to 255 nM for HLA-A*02:01). To ensure constant prediction precision across tumors, we elected to use one allele at a time rather than adapting the predictions alleles to the HLA haplotype of the patient. We selected HLA-A*02:01, –A*11:01, –B*07:02, –B*27:05 and –B*40:01, based on their prior determined in accuracy in predicting nonamer binding affinity^27^ and for their functional diversity. To expedite (repeated) usage of peptide affinity predictions, affinity predictions for all candidate peptides and all HLA alleles encountered in the TCGA patient set were pre-cached in a PostgreSQL database (version 9.5, querying code available in the R package `quickMHC`). Finally, we determined whether T-cell recognition is likely to be prevented by self tolerance. As the majority of mutated antigens derive from single nucleotide variants and, by consequence, are highly similar to their wildtype counterparts, it may be important to exclude candidate peptides that are too similar to the self-ligandome, as thymic negative selection eliminates T-cell clonotypes reactive to antigens from these peptides. To this end, we implemented a ‘self-similarity’ filter^13, 14^ that compares each candidate epitope to a reference list of self-epitopes predicted from the complete human proteome (obtained from UniProt, 2016-10). Candidate peptides were retained when deemed sufficiently different from self, according to the criteria outlined in ‘Similarity to self-repertoire’.

#### Antigen prediction pipeline validation

To evaluate the prediction performance of Neolution when performing predictions either (i) solely based on predicted HLA affinity, or (ii) also incorporating proteasomal processing predictions, or (iii) additionally excluding peptides with similarity to human sequences, a curated list of ‘high confidence’ HIV epitopes (see ‘HIV peptide data for pipeline prediction validation’ and Supplemental Table S2) and ‘lower confidence’ lists of peptides from the IEDB (see ‘IEDB peptide data for pipeline validation’) was processed using it. From these sets of predictions, prediction precision (PPV), sensitivity (TPR) and false positive rate (FPR) were computed using custom R code. To compute 80% confidence intervals (Cis) around PPV-estimates, we employed stratified bootstrapping to ensure constant prevalence of T-cell targeted peptides (and hence their presence) across bootstrapped samples. After quality metrics were computed for bootstrapped samples, the 10^th^ and 90^th^ percentiles of these metrics were taken as the upper and lower boundaries of the reported 80% Cis.

### Somatic variant and neoantigen load tallying

In order to compare the propensity of different classes *c* of a particular classification *C* (e.g., DNA damage types) to yield HLA-A*02:01-antigens, we needed to tally the load of mutations and neoantigens for each class *c* in classification *C*. The majority of mutations can be unambiguously annotated as deriving from a single class *c* (e.g., ‘missense mutation’ in the transcript effect classification). However, some somatic mutations overlapping with the genomic loci of multiple genomic features (i.e., transcripts and genes) have different effects on these genomic features. To allow variants to belong to multiple classes *c* of a classification *C* during the tallying of variants, each variant was partially assigned to a class *c* based on the fraction of transcripts affected by the variant for which the effect of the mutation can be classified as *c*. This way, each variant potentially distributes its contribution among multiple classes, but its total contribution never exceeds unity (1). The total variant load of class *c* in a genome 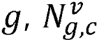, then becomes

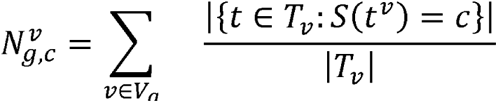

where *v_g_* denotes the complete set of variants in genome *g, T_v_* the set of (Ensembl reference) transcripts affected by variant *v*, the *s* operator returns the variant effect classification of *v* on *t* and |*T_v_*| represents the number of transcripts affected by v (i.e., the cardinality of *T_v_*).

A similar consideration of mixed class membership is made during the tallying of neoantigens that can be contributed by multiple somatic variants of potentially different classes. When tallying the neoantigen loads by different classes *c* of classification *C*, an epitope derived from a transcript *t* was taken to result from *c* proportional to the fraction of total variants overlapping with *t* classified as *c* with respect to *t*. This nuance is necessary as variants may have different effects on different transcripts. Combined, the peptide load contributed by somatic variant class *c* in genome 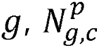, becomes

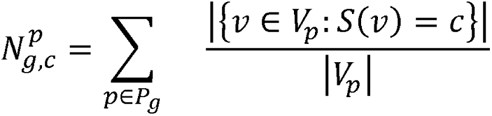

where *P_g_* denotes the complete set of peptides passing all filtering steps for genome *g, v_p_* the set of variants contributing to peptide p, the s operator returns the class of its argument and and |vp| represents the number of variants contributing to peptide *p*.

### The ‘two group’ strategy for testing neoantigen depletion across unpaired samples

As immune pressure against mutations associated with HLA-A*02:01-peptides is not expected in patients that lack the HLA-A*02:01-allele and also any other alleles with similar binding profiles, we could compare the HLA-A*02:01-yield rates in HLA-A*02:01-positive patients (test set) with the yield rates in HLA-A*02:01-like negative patients (reference set). We did this both in a patient specific manner, comparing the distributions of patient specific yield rates against each other, and by aggregating all mutations and neoantigens into a meta-patient before comparing these point estimates. In the former case, distributions were compared using a two-sided Wilcoxon rank sum test implemented in the R base package (`wilcox.test`). In the latter case, proportions of neoantigens over total mutations were compared using a chi-squared test as implemented in R base package (`prop.test`). Effect sizes shown by color in Figures S3C & S3D are log_2_ differences, computed by logging the medians of the yield rates in the test and reference sets and subtracting them from each other. All FDR-multiple testing corrections were done per analysis (i.e., columns in Figure S3D) with Benjamini-Hochberg’s procedure^28^ using the `p.adjust` function in R.

#### Detection power analysis

As mutational load and sample count both influenced the statistical power to detect a signal of epitope loss in our meta-patient immune editing analyses, the limits of detectable immune editing varied between sequencing projects. To determine these detection limits, we first normalized the observed yield rate in the test set to that observed in the reference set, equalizing any potentially pre-existing neoantigen yield rate imbalances between the test and reference sets. This way, we would measure the required rather than additionally required epitope loss for statistical significance. Next, we set out to determine the required immune pressure (*IP*), defined as the fraction of binding peptides that are lost compared to the reference (i.e., the relative yield rate decrease in test set patients) to reach statistical significance. As only true positive predicted binding peptides are immunologically visible and hence targetable, the prediction precision (*PPV*, fraction of true positive and all predicted peptides) downscales the *IP* such that the product of the two determines the actually observed immune strength, *IP_o_*: *IP_o_* = *IP* × *PPV*. As such, we continuously lowered the neoantigen yield rate in the test set by a factor 1-*IP_o_*, increasing *IP_o_* from 0 to 1 in increments of .01, while testing for statistical significance using a chi-squared test for equal proportions (`prop.test` as implemented in R). For each combination of a neoantigen prediction *PPV* (*PPV* ∈{.1,.2,.3,.4,.5,.6}) and a tumor type, the required immune strength to reach this statistically significant effect size was recorded.

### Using the silent mutational load to predict expected neoantigen load and scan for immune editing

Rooney et al.^10^ compared the observed neoantigen load to an expected neoantigen load computed from the silent mutational load, which can be assumed not to be penalized by T-cell pressure as synonymous mutations, unlike non-synonymous mutations, do not form neoantigens. The silent mutational load was subsequently used to estimate exposure to DNA damaging processes and thereby infer the number of (neoantigenic) non-synonymous mutations expected in the case of no selection pressure during tumor outgrowth. Assuming that the ratios between (1) synonymous mutations and non-synonymous mutational loads, and (2) non-synonymous mutations and neoantigen counts are on average equal between tumors, it is possible to compute the expected or predicted number of neoantigens from the silent mutational load and two sequentially applied conversion factors. The expected non-synonymous mutational load *NS_p_* is estimated from the observed silent mutational load by multiplying it with the globally estimated conversion factor *c_s_*⇒*_Ns_*. Next, *NS_p_* is used to compute the expected number of neoantigens *E_p_* by multiplying it with the globally estimated conversion factor *c_NS_*⇒*_E_* We finally end up with *R*, the ratio between observed and predicted neoantigens for an individual sample

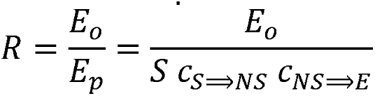

which relates individual samples to the remainder of the cohort which was used to compute the conversion factors.

To model the variable prevalence of various mutational processes operative in the genomes under analysis, Rooney et al. performed this analysis in a mutational spectrum specific fashion, i.e., by taking into consideration the nucleotides directly on the 5’ and 3’ sides of the mutated basepair (e.g., `AC>TG`, a `C>T` mutation flanked by an A and G). Following Rooney et al., conversion factors are here computed specifically for each of the 192 possible spectra. As such, *R* becomes the ratio of the observed neoantigens and the sum of the spectrum specific expected neoantigen loads.

There are a number of differences between the neoantigen prediction strategy employed here and in Rooney et al., precluding a direct copy of their methodology. First, the prediction pipeline employed by Rooney et al. does not account for indels and for interactions between mutations, i.e., potential interactions between mutations in the form of co-occurrence of mutations in a nonamer-spanning genomic sequence. To eliminate this source of possible discrepancy, the *NS* and *S* input data for this analysis are restricted to missense mutations. Analogously, peptides resulting from non-missense mutations are excluded. This means that a neoantigen yielded by for instance a missense mutation and an indel will not increase the missense mutation tally by one half but instead is excluded – it would not have been included in the neoantigen predictions used by Rooney et al.. Second, Rooney et al. performed neoantigen predictions for patient-matched HLA types whereas we elected to use HLA-A`*`02:01-predictions for all analyzed donors because of the superior binding affinity predictions as compared to many other alleles. To further harmonize our methodologies, we excluded all variants (and associated neoantigens) located in driver genes, as these were also excluded by Rooney et al. (driver genes identified using MutSig, Table S6A from Rooney et al.). We performed these analyses on i) either only clonal mutations or all mutations, ii) in an either mutational context specific fashion or not for more robust conversion factor estimates (through including more events per conversion factor), and iii) using our 4-filtering neoantigen prediction pipeline or using predicted HLA affinity only, the latter to be more consistent with Rooney et al.. Conversion factors were recomputed per analysis on all included samples (i.e., pan-cancer).

### The ‘continuous’ detection strategy – correlating neoantigen yield rates to HLA presentation scores across patients

Similar to the discrete-group approach to immunoediting testing, the fundamental idea here is to compare neoantigen yield rates for a particular four-digit HLA class I allele (hereafter called focus allele, e.g., HLA-A*02:01) between tumor samples containing HLA alleles that differ in their ability to present peptides that can be presented by this focus allele, but now analyzed on a continuous scale. In case depletion of neoantigenic non-synonymous mutations does occur, tumor samples that contain HLA alleles that show a large degree of overlap in binding pattern with that of the focus allele (i.e., that have a high presentation score, *h*) should show less neoantigens per mutation (low neoantigen yield rate, *r*) on average, as compared to tumor samples that are poor at presenting these peptides (low *h*). We assess this difference in *r* between tumor samples that are high and low in *h* using linear regression.

#### HLA presentation corroboration

Overlap in HLA presentation (HLA corroboration score, Figure 2A) was assessed through overlap in predicted binding affinity over all 10,072,577 candidate nonameric neo-peptides that we processed for our patient cohort and for all 227 identified HLA class I alleles, in order to select a subset of all 20^9^ potential nonamers that is representative of the human antigenome. The asymmetric HLA corroboration score for alleles *I* to allele *j* was computed as the fraction of peptides binding to *j* that are also predicted to bind *I* and thus falls in the range [0, 1]: 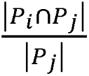, where *P_i_* is defined as the set of peptides presented by HLA allele *i*. Binding peptides were defined as having a NetMHCpan3.0 predicted percentile rank <= 1.9 (which corresponds to 255 nM binding affinity for HLA-A*02:01). Three other thresholds for the definition of binding peptides (1, 3 and 4, corresponding to 100, 500 and 1,000 nM binding affinity for HLA-A*02:01) did not substantially alter our results (data not shown).

**Figure 1:**
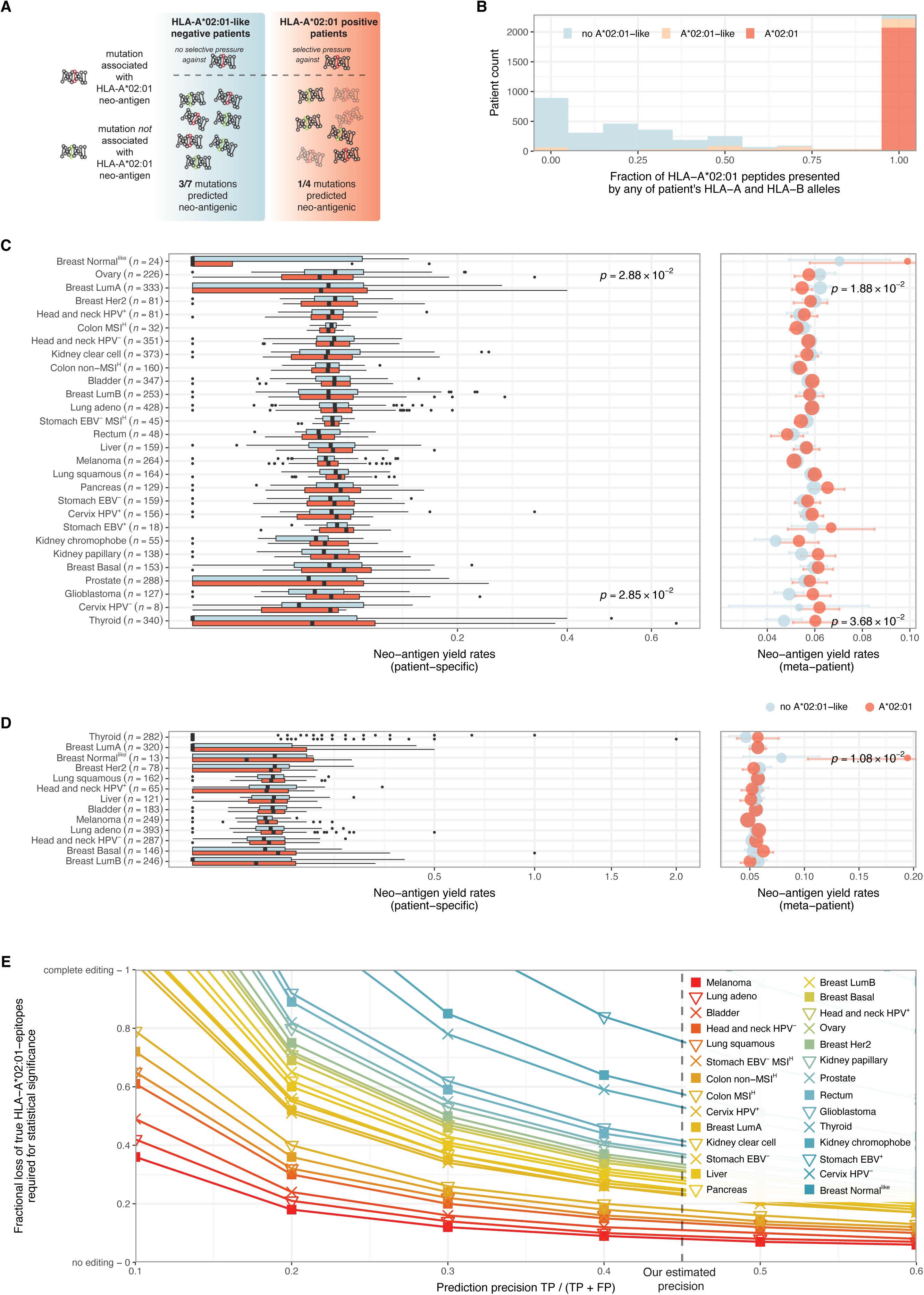
No detectable genomic loss of predicted HLA-A*02:01 neoantigens using a discrete two-group detection strategy. **A**. Schematic overview of analysis strategy. Mutational events that yield HLA-A*02:01-restricted neoantigens can undergo negative selective pressure in HLA-A*02:01-positive patients (the test group), but not in patients that lack this allele as well as other HLA class I alleles with similar peptide binding profiles (“HLA-A*02:01-like negative reference patients”). As a consequence, genomic loss of antigenic HLA-A*02:01 mutations would be reflected in a lower HLA-A*02:01 neoantigen yield rate in HLA-A*02:01-positive patients as compared to HLA-A*02:01-like negative patients (in the indicated example, 3 out of 7 vs. 1 out of 4 non-synonymous mutations yield predicted HLA-A*02:01 neoantigens). **B.** Distribution of the fraction of HLA-A*02:01 binding peptides presented by any of a patient’s HLA class I A and B alleles, according to HLA-A*02:01-allele and HLA-A*02:01-like allele status. Patients in the HLA-A*02:01-like negative reference group present 16.7% (median) of predicted HLA-A*02:01 binding peptides by any of their HLA class I A and B alleles. **C.** Neoantigen yield rates of SNVs in HLA-A*02:01-positive and HLA-A*02:01-like negative tumor samples. Left: distributions of patient specific neoantigen yield rates. Right: ‘meta-patient’ yield rates, in which mutations, and hence neoantigens, are grouped per tumor type. Unadjusted p-values are shown for tumor types where they are smaller than or equal to 0.05. None of the comparisons remained significant after correcting for multiple testing. **D.** As in C., but excluding both subclonal mutations and mutations for which loss may potentially confer a cell intrinsic fitness cost (driver mutations and mutations in essential genes that display LOH). Unadjusted p-values are shown for tumor types where they are smaller than or equal to 0.05. None of these remained significant after correcting for multiple testing. **E.** Power analysis of neoantigen depletion detection strategy. The effect of prediction precision on the fractional loss of predicted neoantigenic mutations required to achieve statistical significance is depicted on a tumor type-specific basis. Dashed vertical line depicts precision of the here employed epitope prediction pipeline, indicating that genomic editing of a minimum of 9% and 18% of true HLA-A*02:01 neoantigens would have been detected in melanoma and colon MSI^H^, respectively, assuming no multiple testing correction of p-values.

**Figure 2.**
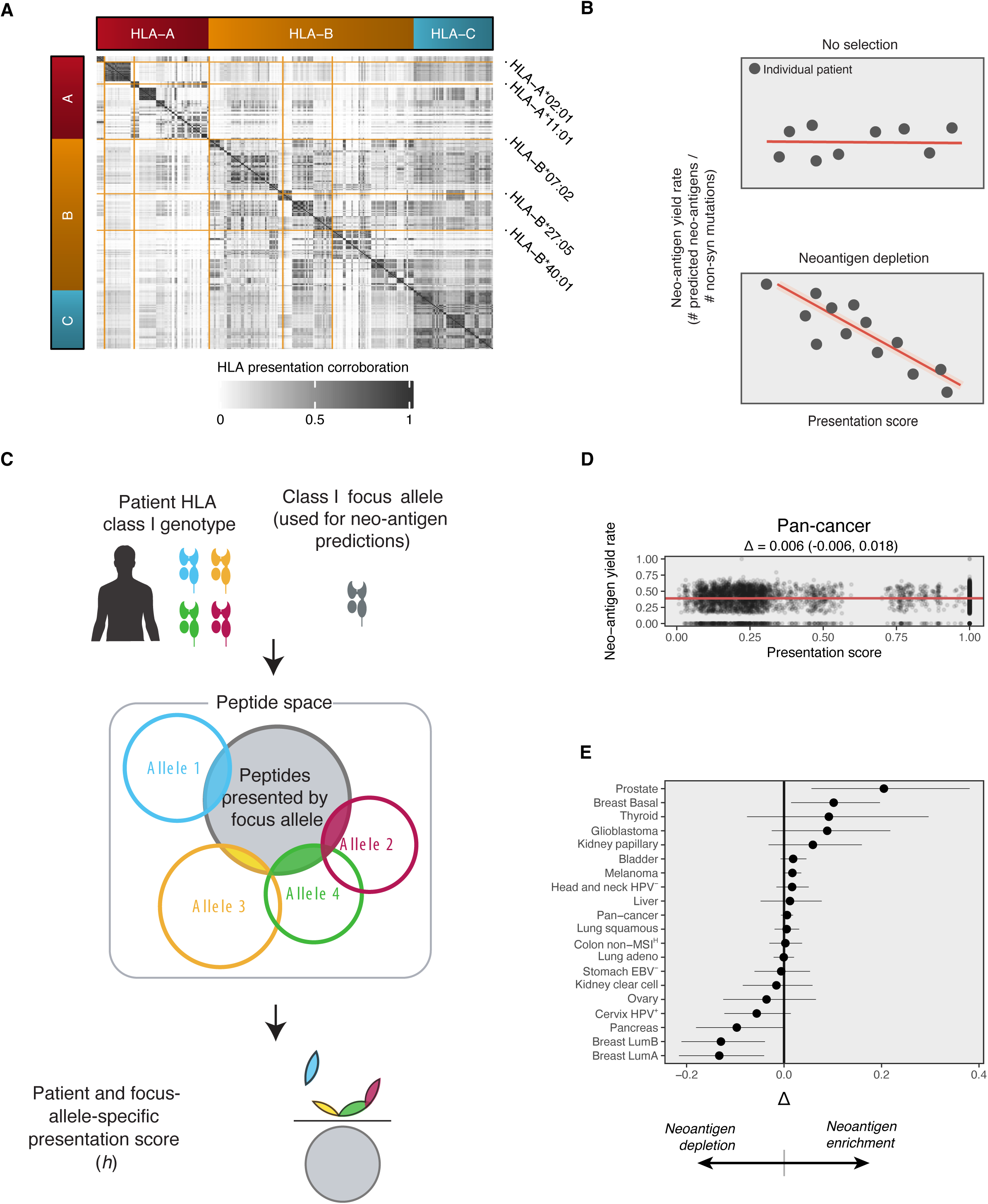
A continuous strategy to detect neoantigen depletion. **A**. Overlap in predicted peptide presentabillity across all detected HLA class I alleles (A, B and C) across the TCGA patient samples included in the analyses. Matrix reflects the fraction of peptides presented by alleles in the rows (i.e., NetMHCpan 3.0 rank percentile <= 1.9), that are also presentable by the alleles in the columns (same presentation criteria). Alleles in both rows and columns are alpha-numerically ordered. The five highlighted HLA-A and –B alleles were selected based on NetMHCpan3.0 prediction accuracy, HLA diversity and to minimize overlap in predicted binding capacity within the group. **B.** Schematic of the continuous (as opposed to two patient-group) methodology to evaluate neoantigen depletion. To account for overlap in (predicted) HLA binding, a continuous score was used that captures a sample’s capability of HLA-presenting peptides that are associated with an HLA allele of interest (the ‘focus allele’). This score is then regressed against the neoantigen yield rate for this focus allele. A negative slope in this regression would indicate that fewer neoantigen-encoding mutations are detectable in samples that have a high capacity to present these. **C.** Schematic of the HLA presentation score (*h*), reflecting the fraction of peptides presentable by the focus allele, that is also presentable by at least one of the HLA alleles carried by a patient. **D.** Pan-cancer result with a basic parameterization of the analysis, as in Figure 1C. **E.** Forest plot of the observed Δ values and associated 80% confidence intervals.

#### Computation of tumor sample and HLA allele-specific HLA presentation scores

For the combination of a the HLA class I repertoire of an individual tumor sample and a particular focus allele for which immunoediting analysis was performed, the HLA presentation score (*h*) was defined as the fraction of peptides that is presentable by the focus allele that is also presentable by one or more of sample’s HLA alleles, i.e., 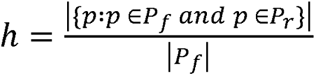, in which *P_f_* is the set of peptides presentable by the focus allele and *P_r_* is the set of peptides presentable by the sample’s class I HLA allele repertoire.

For analyses where allelic loss of HLA alleles (inferred using LOHHLA, see section ‘Annotation of allele-specific HLA loss’) is integrated into *h_i_*, alleles identified to have been lost were excluded from *A* before computation. Allele amplification was ignored. Where the ‘Presentation score’ setting was set to ‘HLA A, B’ (rather than ‘HLA A, B, C’), C-alleles were excluded, regardless of other settings.

#### Mass spec validation of HLA presentation score

To evaluate the validity of the sample-specific HLA presentation scores, we reasoned that mass spec-eluted HLA ligandomes should be enriched for peptides associated with HLA-alleles for which the corresponding patient has a high presentation score. To test this, we obtained mass spectrometry data of HLA-eluted peptides from 26 samples^29^ through the SysteMHC atlas (project ID: SYSMHC00010). We inferred the 4 HLA-A and –B alleles of each sample’s corresponding HLA repertoire by considering the HLA alleles to which peptides had been assigned by the original investigators, assuming homozygosity and two allelic copies when only one HLA-A or B-allele was reported in total. Next, we compiled sample specific reference proteomes by aggregating all source proteins with which the observed peptides were annotated by the original investigators, so as to work around the unavailability of matched RNAseq data for these samples. We then assessed which peptides were expected to be present in the HLA ligandome of each combination of a sample and a focus allele by extracting all 9-mers from these proteins, using the UniProt human reference proteome (UP000005640_9606) as a reference, and annotating these peptides with a predicted affinity for the focus allele. The minority (<.1%) of peptides containing either a U (selenocysteine) or X (unknown amino acid) where excluded a priori. Having filtered these peptides, with passing peptides having an HLA affinity percentile rank <= 1.9, we assessed which fraction of filter-passing peptides was observed in the experimental data to arrive at a quantity we interpreted here as prediction precision. Such predictions precisions were computed for all combinations of all 26 samples and 226 focus alleles (all HLA-A and –B alleles detected in the TCGA patients we analyzed).

#### A continuous approach to testing for immunoediting

We modeled the mean rate at which mutations yield neoantigens for a particular tumor type *r* and how that rate is modulated by the HLA presentation score (*h*) for the focus allele of the analysis. We thus define:

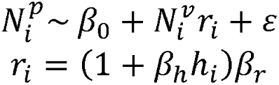

where 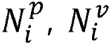 are the neoantigen and mutational load of the patient indexed by *i*, respectively, and *r_i_* is the patient specific neoantigen yield rate, consisting of a global neoantigen yield rate *β_r_*, shared across patients, and a second term that describes the degree to which the HLA presentation capability (presentation score *h_i_*) shifts this global yield rate in a patient-specific manner. This second term consists of the patient-specific presentation score *h_i_* and *β_h_*, the fractional degree to which *r_i_* is modulated by a unit incease in *h_i_*. Finally, *β*_0_ is an intercept term (the predicted number of neoantigens when *N^p^* =0 and *h*= 0) and ε represents the model residuals 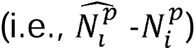.

Integrating these two expressions and removing the coefficients, we get the following R model formula to be used in a R (g)lm:

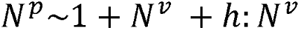

Such that the first coefficient represents an intercept term, the second coefficient represents *β_r_* and the third *β_h_*.

A subanalysis was deemed reliably estimable if the p-value associated with *β_r_* (i.e., the base *r*, the expected neoantigen yield rate for patients with *h*= 0) was < 0.05. In cases of too few samples or the absence of a (linear) pattern in the data, this did not happen.

We were ultimately interested in the mean fractional difference Δ (delta) in neoantigen yield rate between patients with full HLA-presentation capability for the focus allele in question (*h*= 1) and those with no capability whatsoever (*h*= 0), which boils down to the following expression:

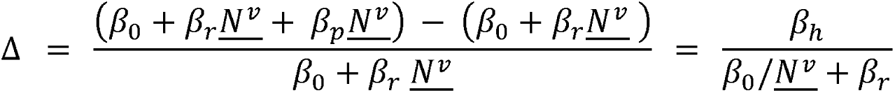

in which *<u>N^v^</u>* is the median number of mutational events across patients. The uncertainty in Δ was estimated using Monte Carlo simulation. We generated 10^4^ random samples from the multivariate normal distribution of the regression coefficients (mean vectors from `coef` and covariance matrix from `vcov` methods of the R fit objects) using the `mvrnorm` function in the R MASS package (version 7.3.54). We computed Δ for each of these simulations and report the 10^th^ and 90^th^ percentiles of the resulting distribution (Figure 2D). Both *N^v^* and *N^p^* were transformed with *f*(*x*) = *log*10(*x* + 1) prior to regression analysis.

#### Permutation testing Δ for significance

We employed permutation testing to assess the statistical significance of observed Δ values. Specifically, the presentation score for each sub-analysis was permuted 250 times and the resulting Δ was computed for each of those permutations. Any downward or upward bias resulting from distributional imbalances and other potential violations of linear model assumptions will be accounted for using this approach. We assessed the fraction of permutation Δ values that was larger than the observed Δ, a quantity we refer to as *q* for quantile. Small values of *q* (< 0.05) are indicative of a substantial deviation from the permutation null distribution and would be consistent with immunoediting. Large values (0.95) would likewise be consistent with neoantigen enrichment.

#### Detecting parameterizations associated with editing

Aggregating all sub-analyses, each a unique combination of all the potential discrete settings to make, resulted in a table with ‘ID’ columns, one for each of the configuration categorical variables, and columns for the resulting observed statistics, Δ and *q*. As we wanted to understand which settings increase the likelihood of observing negative Δ values, we sorted this table for either one of these statistics and univariately tested all ‘ID’ variables independently for enrichment. We repurposed fast preranked gene set enrichment analysis testing here, as implemented in the fgsea R package (package version 1.18.0). For each level of a categorical variable (e.g., ‘A*02:01’ when considering the focus allele) occurring in at least 100 sub-analyses, we ran `fgsea::calcGseaStat` with `scoreType` `=` `‘std’` on the rankings of the sub-analyses and with all the positions of the level in question as the argument to `selectedStats`. The `calcGseaStat` function computes a running-sum statistic and returns the location where that statistic deviates most strongly from that expected under random shuffling of the level: the enrichment score (ES). We report the mean-centered version of this statistic, for each setting separately. Mean values were computed over all enrichment scores and deducted from the original ES.

### Software availability

The TCGA data download is encapsulated in the R package `firehosedownload` (github.com/slagtermaarten/firehosedownload). Neoantigen predictions are performed using our custom software packages `VarContext` (github.com/schumacherlab/varcontext, projecting mutations on transcript RNA sequences to compute candidate neoantigenic peptides), neolution-prep & neolution-live (github.com/schumacherlab/{neolution-prep,neolution-live}, for annotation and filtering of candidate peptides), supported by the `quickMHC` package (github.com/slagtermaarten/quickMHC). Neoantigen tallying is further wrapped in the `fasanalysis` (an acronym for foreign antigen space analysis) package (github.com/slagtermaarten/fasanalysis). Immunoediting analysis code, running the analyses and generating the figures presented in this work, is available at github.com/slagtermaarten/immunoediting. With the exception of VarContext, which is implemented in Perl, all functionality was implemented in R.

## Results

### Design and evaluation of an HLA-I antigen prediction pipeline

To study Darwinian selection against neoantigenic non-synonymous mutations, we first developed an (neo–)epitope prediction pipeline and optimized it with respect to prediction precision and sensitivity. The pipeline annotates candidate nonameric peptides with the output of four tools that jointly model the major requirements for (neo–)antigen presentation: RNA expression of the mutant DNA sequence, predicted proteasomal processing, predicted HLA-binding, and self-similarity of encoded peptides (Figure S1A). To tune the parameterization of this pipeline, we first identified a set of peptides within the 3,094 nonameric peptides present in the HIV genome for which T-cell recognition in the context of the HLA-A*02:01 allele, the most common HLA class I allele in US and European populations, had been demonstrated unambiguously. Specifically, by querying the HIV Molecular Immunology Database^20^, we identified peptides that met the following criteria: (i) HLA subtyping information had demonstrated restriction by the HLA-A*02:01 subtype; (ii) T-cell responses had been observed in at least 3 patients; and (iii) such T-cell responses had been observed in material from HIV-infected patients, rather than induced in vitro or in animal models. This resulted in a set of 32 epitopes in the HIV genome for which presentation by HLA-A*02:01 and recognition by the human T-cell repertoire had unambiguously been established (Table S2). Subsequently, we used this epitope set to compare the sensitivity and precision of epitope prediction strategies that either solely relied on predicted HLA binding affinity^21^, or that also integrated filters that predict proteasomal processing^22^ and similarity of a candidate epitope to self-peptides predicted from the human proteome^23^, STAR Methods). As compared to epitope predictions solely based on predicted HLA affinity, we found both incorporation of proteasomal processing and exclusion of peptides with similarity to human sequences to improve prediction precision (Figure S1B), in line with other work^24^. Selection of an affinity threshold of 255 nM yielded a good balance between precision and sensitivity (Figure S1B-D) of this prediction pipeline. We note that experimental data on T-cell recognized HLA-A*02:01 peptides in the HIV genome may still be incomplete, rendering these performance estimates lower bounds on true prediction performance.

To complement the HIV-pipeline validation, we did a similar validation on peptides derived from the IEDB for 10 viruses. The coverage of T-cell recognized peptides in the IEDB database was likely substantially lower, as in this database we retrieved only 9 out of the 33 T-cell targeted HIV-peptides that were identified in the Los Alamos database, and this probably underlies the observation that much stricter pMHC affinity-filtering yielded optimal precision for the IEDB derived epitope set (Figure S1E,F).

### The two patient-group methodology

To next determine whether neoantigen depletion detectably shapes the mutational landscape of human cancer during tumorigenesis, we first compared the number of predicted HLA-A*02:01 neoantigens per mutational event (the neoantigen yield rate, *r*) for tumors of HLA-A*02:01-positive patients (test set) and tumors from patients that lack the HLA-A*02:01 allele and also other HLA class I alleles with similar peptide binding profiles (reference set, see below for the filtering of HLA alleles with similar peptide binding profiles). As the latter patient group cannot present these predicted HLA-A*02:01-restricted neoantigens, this group provides a reference that can be used to calculate to what extent T-cell pressure has shaped the repertoire of neoantigens as predicted on tumor mutation data (Figure 1A). In addition, by focusing on neoantigen predictions for a single class I HLA-allele, rather than the diverse set of alleles carried by any individual patient, an equal and, in case of HLA-A*02:01, high prediction accuracy is guaranteed across patients.

To be able to assign patients to either the test or reference groups, we first assessed the similarity in peptide binding properties between HLA-A*02:01 and all other HLA class I alleles encountered within our patient set, by computing for each observed HLA-allele the fraction of the 62,833 predicted HLA-A*02:01 neoantigens observed in this cohort that it was also predicted to present (corroboration index, Figure S3A). HLA class I alleles that were predicted to present more than 20% of the set of HLA-A*02:01-peptides were classified as ‘HLA-A*02:01-like’ (37 of 224 HLA class I alleles). Subsequently, patient samples expressing at least one such allele were removed from the analysis, resulting in an HLA-A*02:01-positive test group of 2,345 patients and a reference group of 2,628 patients that lacked HLA-A*02:01 and also any HLA class I allele with substantially overlapping binding properties. Residual binding repertoire overlap between the groups was small, with reference group patients being able to present 16.7% (median) of their predicted HLA*02:01 presented peptides using any of the up to 6 HLA class I alleles that they expressed (Figure 1B), i.e., 6-fold lower than in the HLA-A*02:01-positive test group.

To test for preferential loss of mutations that yield predicted HLA-A*02:01-presented neoantigens, we compared *r* between test and reference patient groups per tumor type. This analysis revealed a significantly lower *r* in the test set for ovarian carcinoma, while a higher *r* in the test set, i.e., a presumed neoantigen enrichment, was observed for glioblastoma. However, neither of these results remained significant after multiple testing correction (Figure 1C, left), indicating a lack of detectable neoantigen depletion above the noise levels in the data. Absence of significant depletion was also observed when aggregating all mutations for each tumor type into one HLA-A*02:01^+^ and one HLA-A*02:01-negative ‘meta-patient’, an approach that is expected to increase analysis robustness in case of low mutation numbers (Figure 1C, right).

A potential limitation of this analysis strategy could be formed by the presentation of a fraction of HLA-A*02:01-presentable peptides by the aggregate of all the other HLA-alleles in the reference set patients, such that a degree of ‘background’ immune pressure can be expected to occur against HLA-A*02:01-peptides. To test robustness with respect to the stringency used to create the non-HLA*A-02:01-like reference patient set, we performed meta-patient tests using an increasingly strict HLA-similarity threshold (i.e., allowing a progressively lower fraction of HLA-A*02:01-presented peptides to be presented by the non-HLA*A-02:01-like reference patient set). Using this approach, no systematic increase in statistical significance was observed (Figure S3B), suggesting that the (low-level) overlap in peptide HLA-binding profiles was unlikely to confound this analysis.

As the probability of epitope presentation depends on the RNA expression level of the associated gene^25–28^, an increase in RNA expression thresholds for neoantigen predictions may be expected to increase precision (i.e., increase the fraction of truly presented peptides amongst predicted peptides), and could thereby potentially reveal a weak neoantigen depletion signal. Similarly, loss of neoantigens could be postulated to be more apparent among high affinity HLA ligands that are more likely to yield strong T cell targets. To test both possibilities, we titrated both the RNA expression and HLA affinity thresholds of the neoantigen prediction pipeline and re-evaluated HLA-A*02:01 *r* in the test and reference patient sets. Small differences between these two sets were observed when performing pipeline stringency titrations and we did observe significantly lowered *r* for multiple analyses (i.e., rectum, breast LumB, melanoma, head and neck HPV^-^, Figure S3C). Importantly however, a similar number of tumor types with a heightened *r* was once again observed (Figure S3C). We conclude that absence of detectable neoantigen depletion is robust to different configurations of the neoantigen prediction pipeline.

To screen for other factors that could have obscured a weak neoantigen depletion signal, we performed additional analyses in which we excluded mutations for which negative selection could have been counter-acted by positive selection. Specifically, T cell-mediated depletion of oncogenic mutations in driver genes may be expected to be counteracted by the positive effect of these mutations on cellular fitness^29^. Similarly, essential genes that encode neoantigens and of which the wild type copy has been lost (so called essential passengers^29^) cannot be lost without loss of cell viability^29, 30^. Finally, we performed analyses that excluded subclonal mutations, as these could be postulated to have emerged too recently in order for the immune system to have affected their presence. When excluding the aforementioned mutation classes, we again discerned no statistically significant neoantigen depletion, neither when analyzing patient-specific nor meta-patient neoantigen yield rates (Figure 1D, left and right, respectively, fewer tumor types due to lower availability of required data for clonality calling, Methods).

In prior work, the HLA class I-restricted presentation of neoantigens has been reported to shape the repertoire of oncogenic mutations, with individual driver mutations being reported to occur more frequently in patients with HLA repertoires that are less likely to present the resulting mutant peptides^13^. We attempted to validate this observation using our epitope prediction and analysis strategy. Restricting our analysis to the recurrent driver single nucleotide variants (SNVs)^13^, we also did not observe significant neoantigen depletion (Figure S3D). Finally, escape from immune pressure can occur through a variety of genetic alterations, including mutations in components of the antigen presentation machinery, and the loss of T-cell-recognized neoantigens can reasonably be expected to no longer provide a fitness advantage in tumors that harbor such alterations. To restrict our analysis to tumor samples for which no evidence of other known escape mechanisms was present, we excluded all tumors with one or more non-silent mutations in any of 515 genes implicated in resistance to T-cell killing through CRISPRi screening^31^. Application of this tumor sample filtering to all of the analyses reported above resulted in larger differences for at least one tumor type per analysis. However, this did not result in an increased number of tumor types for which significant neoantigen depletion was observed after correcting for multiple testing (Figure S3D). Our recurrent finding of absence of significant neoantigen depletion in treatment-naïve tumors contrasts with an earlier assessment of neoantigen depletion on TCGA data^11^. Adapting this prior analysis strategy to our HLA-A*02:01-centered neoantigen predictions (Methods) again did not reveal any signs of genomic epitope loss (Figure S3E), even when restricting this analysis to clonal mutations (Figure S3F), or when exhaustively testing other possible variants of this analysis (Figure S3D).

Even when using optimized epitope prediction pipelines, neoantigen predictions still contain a sizeable fraction of false positive peptides. As only true positive peptides (i.e., predicted peptides that are true HLA ligands) can potentially be seen by T cells, and hence can be subject to Darwinian selection, prediction precision defines the lower limit for the degree of neoantigen depletion that would be detectable. We determined what degree of neoantigen depletion would have been detectable using the two-group methodology, by computing the required effect size (i.e., fractional loss of true epitopes in HLA-A*02:01-positive tumors) to reach statistical significance given the observed noise in the data. At our estimated neoantigen prediction precision of 0.45 (Figure S1B,C), unadjusted *p*-values of 0.05 would have been reached upon loss of 8-18% of truly presented epitopes in melanoma, lung and colon cancers (Figure 1E), indicating that the true neoantigen depletion signal must have been smaller than that for it to have not been detected.

### A continuous version of the neoantigen depletion analysis

Given the overlap in binding profiles of different HLA class I alleles (Figure 2A), and the resulting continuous rather than bi-modal distribution of HLA-presentation overlap with HLA-A*02:01 across samples (or any other HLA-allele, hereafter called the focus allele), a statistically more powerful approach could be to test for a (negative) association between focus allele-presentation capability and neoantigen yield rate (*r*) across samples. Modelling focus-allele presentation capability with a quantity we call the HLA presentation score (*h*, methods), a detectable signal of neoantigen depletion (i.e., depletion of neoantigenic, non-synonymous mutations), should on average lead to a lower *r* in samples with a high *h*. That is, a linear regression between *h* and *r* should yield a negative slope (Figure 2B, bottom). In contrast, a slope of zero would indicate HLA presentability not to be associated with depletion of mutations carrying predicted neoantigens (Figure 2B, top). We modeled *h* as the fraction of unique, theoretically presentable focus allele peptides that are also presentable by one or more of the patient’s class I alleles (Figure 2C). In this way, *h* will be 1 for patients that do carry the focus allele while those that do not will have values ranging from 0 to 1.

To first evaluate whether *h* models peptide presentation capacity, we analyzed mass spectrometry data of HLA-eluted peptides^32^. Specifically, having inferred the HLA class I repertoire of each patient in this data set, we computed *h* for each patient and potential focus allele combination. Next, for all 9-mer peptides of the expressed human proteome we predicted whether mass spec detection would have been expected solely based on peptide affinity predictions. In case *h* models peptide presentation correctly, the proportion of predicted peptides for a given focus allele to be detected by mass-spec should be correlated to *h*. Confirming our expectation, we observed positive correlations between *h* and the number of detected over expected peptides for 23 of 27 analyzed samples (Figure S4). Having computed *h* across all five focus alleles and all samples, we observed a near-zero correlation between focus alleles (Figure S5A,B). This indicates that analyses using different focus alleles form largely independent and complementary tests within a fixed set of tumor samples, and can hence be seen as independent validations.

We next assessed the relationship between *h* and SNV *r* across all evaluated patients (i.e., pan-cancer) using HLA*A-02:01 as the focus allele, observing a virtually flat, non-statistically significant, slope. To put this slope into perspective, we defined *L* (delta) as the relative difference in *r* between patients at *h*= 0 (no ability to present focus-allele peptides) and *h*= 1 (full ability to HLA-present focus-allele peptides, Methods), and observed a L of 0.006 (80% CI: [-0.006, 0.018], i.e., non-significant enrichment rather than depletion of predicted neoantigenic mutations, Figure 2D). Testing of individual tumor types in this manner showed apparent neoantigen depletion in certain tumor types (Δ< 0), but just as many other types showed a similarly strong apparent enrichment for neoantigens (Δ> 0), likely reflecting noise in the data (Figure 2E).

Analogous to the two-group based neoantigen depletion methodology described above, we next systematically varied all possible settings of the neoantigen prediction pipeline and the continuous neoantigen depletion analysis strategy in order to test the robustness of these results. As the optimal neoantigen prediction pipeline configuration remains uncertain, despite our efforts to determine it (Figure S1B-F), we evaluated all outcomes while varying three settings of the neoantigen prediction pipeline: i) the *HLA affinity rank percentile threshold* that denotes the predicted HLA affinity candidate neoantigens had to reach for predicted HLA-presentation (4 levels, ranging from lenient to highly stringent), ii) *RNA expression*, either evaluated at the gene level or at the variant level (the latter to be sensitive to potential epigenetic silencing of neoantigenic mutations) and iii) the *similarity-to-self* filter that aims to model thymic selection of T-cell reactivity.

We also varied 6 settings that determine how the neoantigen depletion test is performed, independent from the neoantigen prediction pipeline configuration: i) *Variant selection* determines the set of somatic variants based on which *r* was evaluated and included the following classes: all (SNV) genomic variants; frameshifting indels as these may form richer sources of neoantigens that are typically less self-similar^33, 34^ and could thereby be postulated to experience stronger negative selection; only clonal SNVs mutations; SNVs with driver and essential passenger mutations removed^29^; only highly-recurrent driver SNVs, as defined in a prior work^13^. ii) *Focus-allele* reflects the HLA-allele for which both h and r was evaluated (the two main variables of the regression). The focus allele was varied between five HLA class I-alleles that we picked to cover a broad range of HLA supertypes and that showed a relatively high prediction accuracy using NetMHCpan3.0^21^. iii) *LOH in HLA* determines whether allelic loss of HLA class I^35^ was reflected in the presentation score (*h*), i.e., whether HLA alleles that were reliably found to have been genomically deleted were or were not excluded when computing *h*.

We discerned between a high-confidence variant of this variable, in which only patients for which all alleles could be reliably were included in the final regression analysis (“strict LOH HLA”, Figure S6, Methods), and a more lenient version in which any allele reliably found to be lost was excluded from the HLA presentation score alleles, independent of the assessment quality for other alleles in the same sample (Methods). iv) *C-allele in presentation score* reflects whether both HLA-A, –B, and –C alleles, or only the HLA-A and –B alleles, were included when computing *h*. As most known neoantigens are HLA-A or HLA-B restricted^36^ and HLA-C might be expressed at a lower level^37^, exclusion of HLA-C could lead to more accurate estimates. v) *T cell-resistance*, entails the removal of patient samples from the analysis that carry one or multiple mutations, other than genomic loss of HLA class I loci, in immune evasion genes as identified using CRISPRi-screening^31^, in order to exclude samples in which immune pressure may be reduced by an independent genetic event. Here, patient samples were excluded using three levels of stringency, reflecting false discovery rate thresholds for genes in the CRISPR-screen of 0.0001 (stringent on genes and hence lenient on sample inclusion), 0.1 (moderate on sample inclusion) and 0.01 (stringent on sample inclusion). vi) *Cytolytic score* determines whether sample inclusion is restricted to samples with a high T cell infiltration, as neoantigen depletion might be more apparent in immune infiltrated tumors^17^. here we restricted the tested samples to those high (>=75^th^ quantile within the tumor type) in cytolytic score^11^ (a transcriptomic proxy for T/NK-cell activity) and those lower in cytolytic score (<75^th^ quantile) to contrast the former results with.

Evaluating all combinations of the settings listed above, we frequently encountered patient subsets that became prohibitively small for regression analysis (i.e., uncertain or no regression coefficients) and neoantigen yield rates that became prohibitively low (resulting in a majority of patients with 0 predicted neoantigens). Restricting ourselves to combinations (so called ‘sub-analyses’) for which i) at least 25 samples had *h* in the range [0, .25] and in the range [0.75, 1], ii) the baseline *r*, i.e., *r* for samples with *h*= 0, could be reliably estimated by the model (methods), and iii) at least 100 samples had non-zero neoantigen loads, left 226,376 of 2,822,400 theoretically possible sub-analyses (8.0%, Figure 3A). As expected, tumor types with a large number of samples allowed more restrictive sample filtering and those with high TMB allowed more stringent neoantigen filtering. As such, individual levels of settings were present with variable frequency between tumor types, such that tumor types could not directly be compared from the set of all filtered sub-analyses (Figure 3B). Assessing Δ across the filtered sub-analyses, we did discern tumor types for which the distribution appeared (strongly) skewed to the negative side (Figure 3C, consistent with neoantigen depletion). However, this distribution simultaneously also appeared to be positively skewed for other tumor types. To assess the statistical significance of these and other sub-analyses, we employed permutation tests. Specifically, for each sub-analysis, we permuted *h*-scores across individual samples in the source data 250 times, evaluated *Δ* for each of these permutations, and assessed the fraction of permutation *Δ*s that was smaller than the original *Δ*, arriving at a quantity we call *q* (for quantile). As *q* virtually never reached below .05 (*n* = 4, 0.002%) or above .95 (*n* = 11, 0.005%, Figure 3C, vertical axis), shifts towards either negative or positive *Δ* (i.e., neoantigen depletion and neoantigen enrichment) did not appear statistically significant.

**Figure 3.**
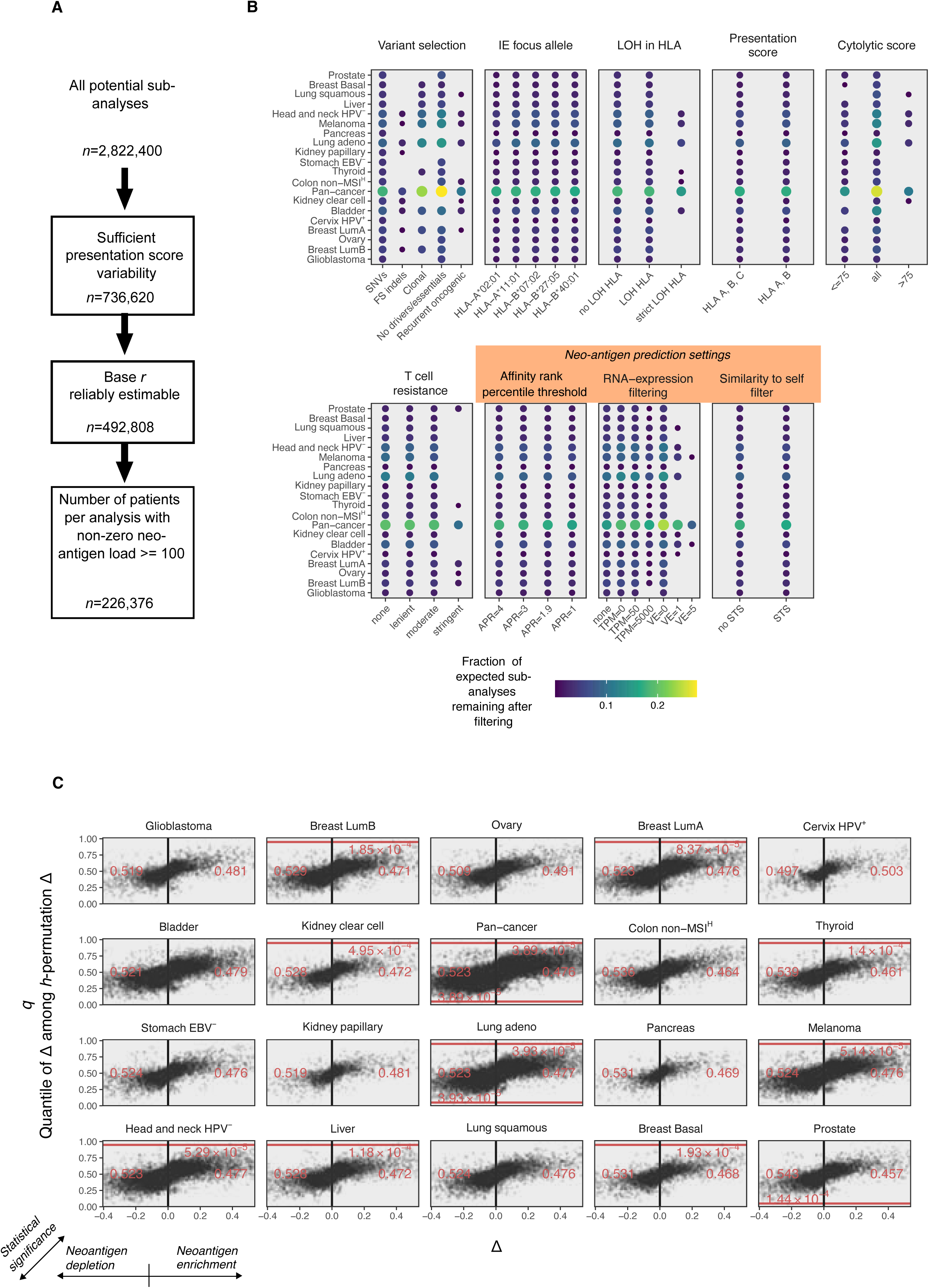
Extended search for neoantigen depletion using the continuous detection methodology. **A**. Filtering of sub-analyses to enrich for sub-analyses with acceptable inferential quality. Blocks denote filtering steps, reported numbers are the number of remaining sub-analyses after filtering. **B.** Sub-analysis composition differs between tumor types because of technical (e.g., the number of patient samples, the quality of DNA/RNA-sequencing data) and biological (mutational load) reasons. Chart shows the fraction of times each factor level (horizontal axis) is observed among the filtered sub-analyses. Levels that lead to strong reductions in predicted neoantigen counts or severely limit sample numbers are only feasible for high mutational load/ more highly represented tumor types. **C.** Volcano-like plot of all filtered sub-analyses, each dot represents a single sub-analysis. Due to compositional differences (see panel B), these plots cannot be used to directly compare tumor types to each other. In case of presence of eccentric sub-analyses (*q* < 0.05 or *q* > 0.95), red lines highlight these thresholds and the fraction of sub-analyses surpassing these are indicated in red.

To characterize these results and identify which individual settings most strongly enriched for neoantigen depletion, we ordered all sub-analyses by *q* and used preranked gene set enrichment analysis^38^ to identify sub-analysis settings that were associated with extreme values in *q* in a univariate manner. As all of the settings are categorical, we compared individual levels (e.g., HLA-A*02:01 for the focus allele) against the combination of all other levels. In this, we recorded the normalized location (range: [-1, 1]) where the rank sum statistic deviated most strongly from that expected under random ordering (i.e., the enrichment statistic^38^). Using this strategy, we did not find evidence that more stringent (and precise) neoantigen prediction could reveal neoantigen depletion (Figure 4A). Surprisingly, repeating this analysis but sorted by *Δ* rather than *q* showed that stringent RNA expression filtering on the variant level on average led to a negative *Δ* for a majority of tumor types (Figure S7A). This was however likely caused by the combination of a heterogeneous distribution of *h* combined with low overall *r* (Supplemental Note 1), and hence artefactual.

**Figure 4.**
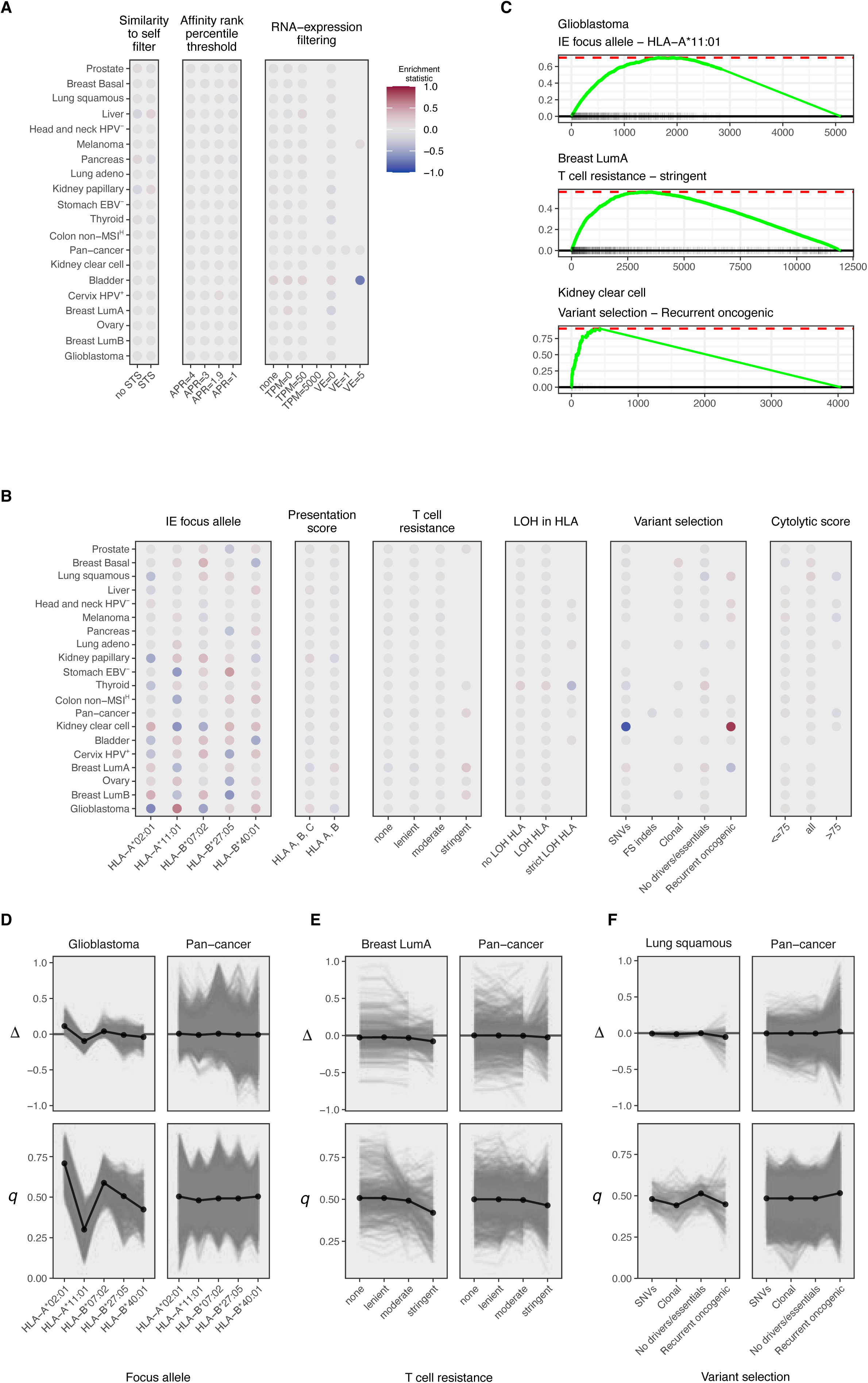
Extended analysis of continuous methodology to assess neoantigen depletion. **A.** Enrichment scores for sub-analysis settings associated with neoantigen prediction methodology, after ranking the sub-analyses by *q*. Note that none of the settings is strongly associated with *q*. ID-variable levels appearing in less than 10 sub-analyses were excluded from analysis. **B.** As in A. but for settings associated with the manner in which neoantigen depletion analysis is performed. As the choice for a focus allele is arbitrary, it is not expected to affect the resulting statistics in case of a detectable and true signal, but it is shown here to be the dominant source of variation. Color scale as in B. **C.** Highlighted individual enrichment-analyses from B, selected for their relatively strong enrichment scores. **D.** Randomly selected groups of identically parameterized sub-analyses, differing only in focus allele, to directly visualize the average effect of variation in focus allele. Glioblastoma and pan-cancer analyses are highlighted to illustrate the relatively strong and weak enrichment scores, respectively, for these tumor types in panel C. **E.** As in D., but varying the subselection of potentially T-cell resistant samples on the horizontal axis. ‘Stringent’ denotes the most rigorous filtering of tumor samples based on the presence of potential immune evasion mechanisms, selecting tumor samples with minimal numbers of mutations that have been associated with immune evasion. **F.** As in D., but varying the somatic variant set on which *r* is evaluated. Frameshift indels were excluded as they could only be reliably assessed on the pan-cancer level (in which they weren’t associated with *q* or Δ).

When evaluating all settings of the neoantigen depletion testing procedure, we observed that the focus allele most strongly affected both *q* (Figure 4B-D) and *Δ* (Figure S7D). Varying the choice of the focus allele allows for semi-independent replication of a neoantigen depletion test, as there is no expected link between the arbitrary choice of an HLA class I allele as the focus allele and the occurrence of neoantigen depletion or enrichment. Importantly, the large observed variation between sub-analyses carried out in this manner and that are otherwise identically parameterized suggests that most of the signal observed can be ascribed to measurement noise. By the same token, we found rigorous exclusion of T cell-resistant tumors to shift *q* towards neoantigen depletion in a slight but consistent manner (5/5 evaluable tumor types, Figure 4B,C,E), but for only 1 out of 5 tumor types (Breast LumA, Figure S8A) this effect was somewhat consistent across the 5 evaluated focus alleles, again suggesting this effect to be spurious. Finally, evaluating *r* strictly on a set of 1018 recurrent oncogenic mutations that have previously been reported to occur preferentially in patients that express HLA class I alleles that are predicted to present these relatively poorly^13^, also might weakly enrich for a signal of neoantigen depletion (7/9 evaluable tumor types, Figure 4B,C,F), but this did not hold up at the pan-cancer level and – importantly – was highly inconsistent between focus alleles for all these 9 tumor types (Figure S8B), again indicating spurious associations.

## Discussion

Using two different methodologies to estimate neoantigen depletion, we observed little if any detectable signal across unmatched tumor samples of treatment-naïve patients, also when controlling for a number of potentially confounding factors. This finding contrasts with part of the existing literature on the topic, in which evidence for neoantigen depletion in treatment naïve tumors was reported. In cases in which we tried to adapt existing methodologies to ours, we also did not detect neoantigen depletion or only observed minimal, non-statistically significant, trends.

It is important to emphasize that we do not see our data as evidence that neoantigen depletion does not occur. Specifically, given the support for immunosurveillance of nascent tumors^39^ and the strong evidence favoring a role for T cell recognition of mutation-induced neoantigens in tumor control^40, 41^, we deem positive selection of tumor cell clones that never acquired or that lost T cell-recognized neoantigens plausible a priori. If neoantigen depletion does indeed occur, how can this be reconciled with the observation that neoantigen depletion is not observable in the genomic analyses presented here? First, our data are consistent with the possibility that a small subset of neoantigens in pre-treatment tumors is lost due to immune pressure, below the level that surpasses the large prediction noise present in our analyses but also in the other approaches that have been explored. Specifically, experimental data suggest that only a small minority of predicted neoantigens (approximately 1%) induces detectable T cell responses^8, 42, 43^. If only this small minority of T cell-recognized neoantigens is at risk of deletion, the maximal depletion signal that could be expected would be proportionally smaller, and would be difficult to detect in the background of predicted epitopes that never led to T cell pressure. On a related note, it is possible that neoantigen depletion does occur at the genomic level, but only or predominantly in later disease stages, or post-(immuno)therapy. Third, neoantigen depletion may be infrequent due to the presence of other, perhaps more potent or more easily accessible, mechanisms of immune evasion^35, 44^. Notably, across tumor types, the median fraction of patients harboring any form of genetic immune escape other than neoantigen loss in their primary tumors was reported to be 0.20 (for metastatic tumors: 0.27) and as high as 0.74 for kidney chromophobe cancer^44^.

To increase the sensitivity of future analyses of genetic information on compendia of tumor samples, it will likely be critical to account for differences in immunogenicity between predicted neoantigens, rather than merely filtering mutations for those expected to be neoantigenic and implicitly assuming equal immunogenicity. At present, technologies to predict the development of T cell responses against a collection of predicted HLA-presented neoantigens are still limited in reliability, despite substantial efforts^45^. Furthermore, if part of this process is stochastic, for instance governed by the occurrence of a specific TCR recombination, the development of accurate predictors of immunogenicity may be challenging. A more attractive approach may thus be to identify very large numbers of T cell recognized neoantigens in prospective clinical studies and to use such epitopes as a wet lab-validated starting point. High throughput T cell repertoire sequencing of T cells with tumor-reactivity signature^46^, along with advances in experimental approaches to screen the reactivity of many T cell clonotypes in parallel, is expected to enable this effort^47–51^.

## Acknowledgments

This work was supported by H2020 grant APERIM (grant agreement number: 633592), Dutch Cancer Society grant NKI 2012-5463 and the Queen Wilhelmina Cancer Research Award. We thank the Research High Performance Computing facility of the Netherlands Cancer Institute for maintaining the computational infrastructure this work was performed on. We thank the patients of the TCGA Program for sharing their samples and the program itself for collecting and making available all source data used herein.

## Author contributions

M.S. designed and performed all analyses unless otherwise stated below. L.F. set up the employed neoantigen prediction pipeline. M.M.v.B. compiled the validation HIV-peptides set and performed informative pilot studies on neoantigen depletion. A.V. implemented VarContext. G.B. performed PAM50 classification of breast cancer samples. T.N.S. conceived and supervised the study. L.F.A.W. supervised the study. M.S., L.F.A.W., and T.N.S. wrote the manuscript. All authors approved the manuscript.

## Data availability statement

The data that support the findings in this study all derive from the TCGA Project. Specific locations of online access are disclosed in the Materials & Methods section.

## Figure captions

**Figure S1.**
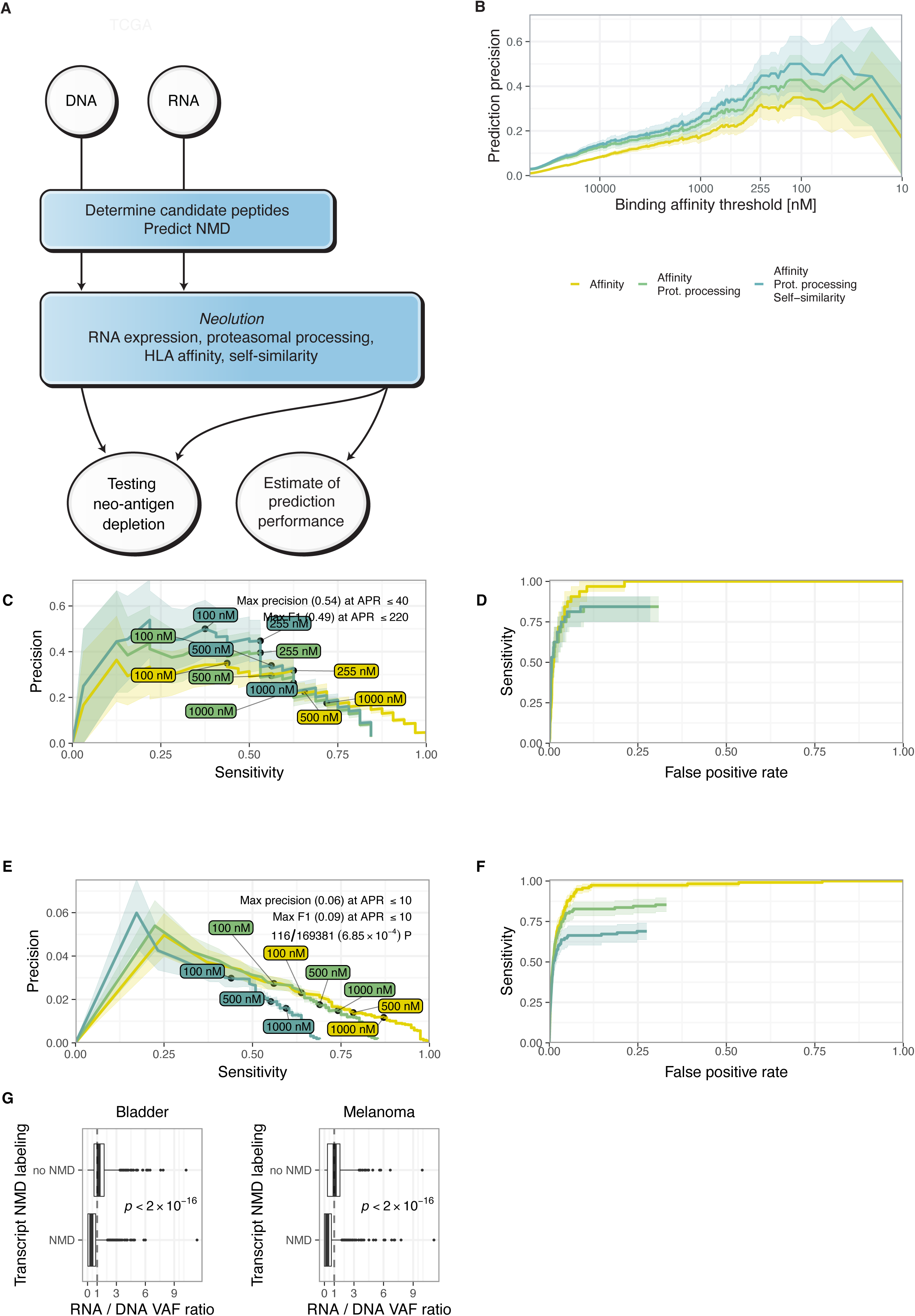
related to Figure 1: Prediction performance for (neo–)antigens and NMD-filter. **A.** Schematic overview of data flow in this work. **B.** Prediction precision as a function of the HLA-affinity filter threshold in the curated HIV epitope set for HLA-A*02:01. Shaded areas denote 80% confidence intervals acquired using stratified bootstrapping. **C.** Parametric plot showing the trade-off between prediction precision (true binders amongst predicted binders) and prediction sensitivity (predicted true binders of all true binders) as a function of the HLA-affinity filter threshold in the curated HIV epitope set for HLA-A*02:01. Increasing the stringency of the affinity threshold increases precision with a trade-off in sensitivity. Dashed red line indicates the 255 nM affinity threshold used for further analyses. **D.** As in A., but plotting the trade-off between sensitivity and drop-out (FPR). Inset: zoom in on the horizontal range of [0, 0.03] of the ROC curve. **E.** As in A., but for the aggregate of all 10 viral genomes in IEDB (Methods). **F.** As in B., but for the aggregate of all 10 viral genomes in IEDB.

**Figure S2.**
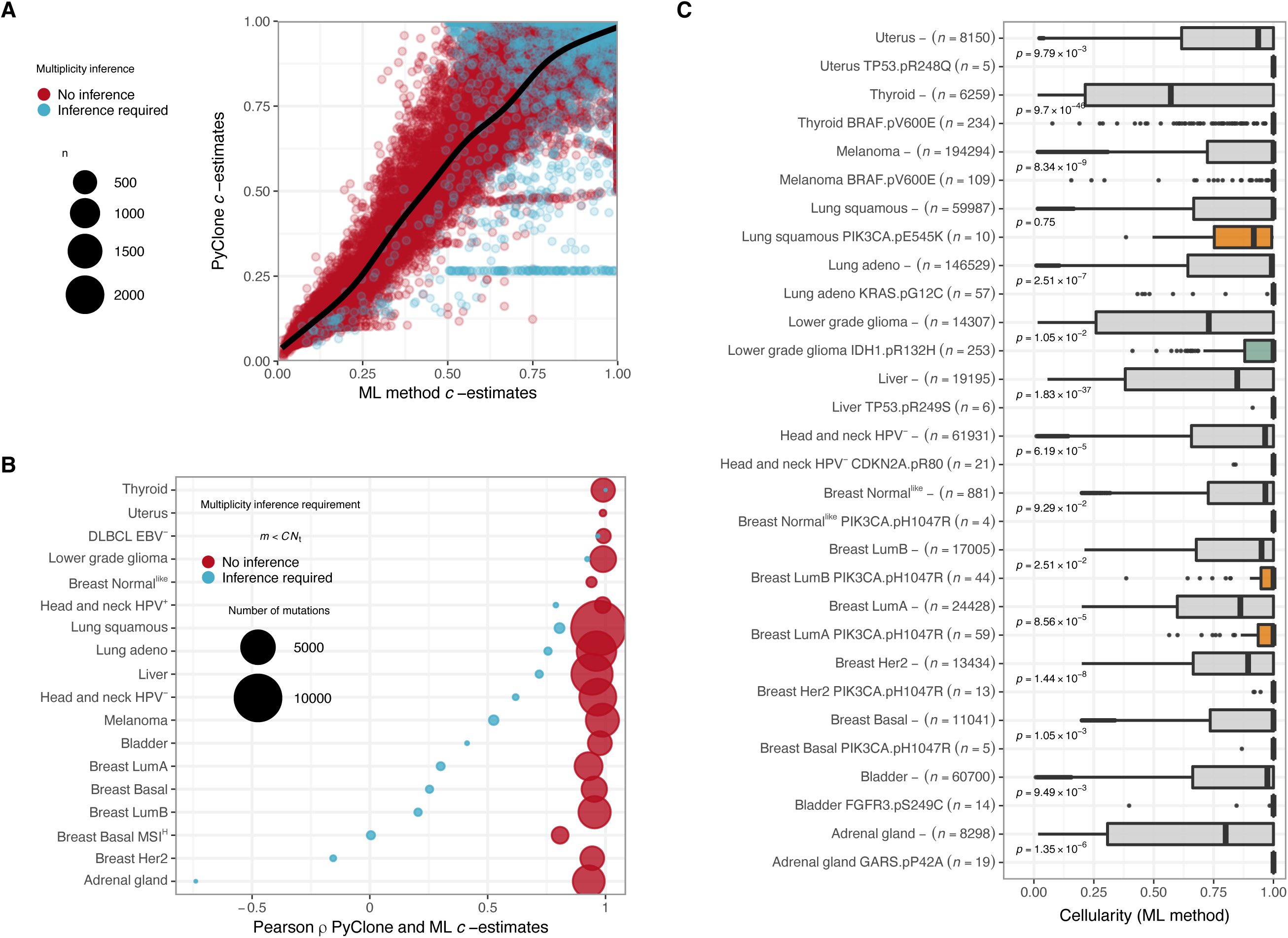
related to Figure 1: Validation of mutation clonality estimates. **A**. Correlation of cellularity estimates obtained by PyClone and maximum likelihood method. Color indicates whether a mutation is located in an amplified region and thus whether multiplicity inference is required. 98.75% of mutations do not strictly require multiplicity inference (blue dots). **B.** Pearson correlation between maximum likelihood and PyClone estimates of cellularity for each tumor type individually. Correlations are high for mutations where the multiplicity is certain but less for mutations for which multiplicity estimates are required (STAR methods). Dot size reflects the number of mutations on which the correlation coefficient is based. **C.** The mostly highly recurrent somatic mutations per cancer types, suggestive of a role in oncogenesis, are significantly more clonal than the aggregate of all non-recurrent mutations (indicated by a hyphen, grey boxplots) for 12/13 evaluated tumor types. **D.** Mutations were grouped based on their effects at the amino acid level. *p*-values reflect one-sided Wilcoxon rank sum tests between non-recurrent and the most highly recurrent (across included tumor samples) mutations. Esophagus cancer and DLBCL were left out of this comparison as the most recurrent mutations were positioned in non-oncogenes for these tumor types (*CEP170* and *MUC6*, respectively).

**Figure S3.**
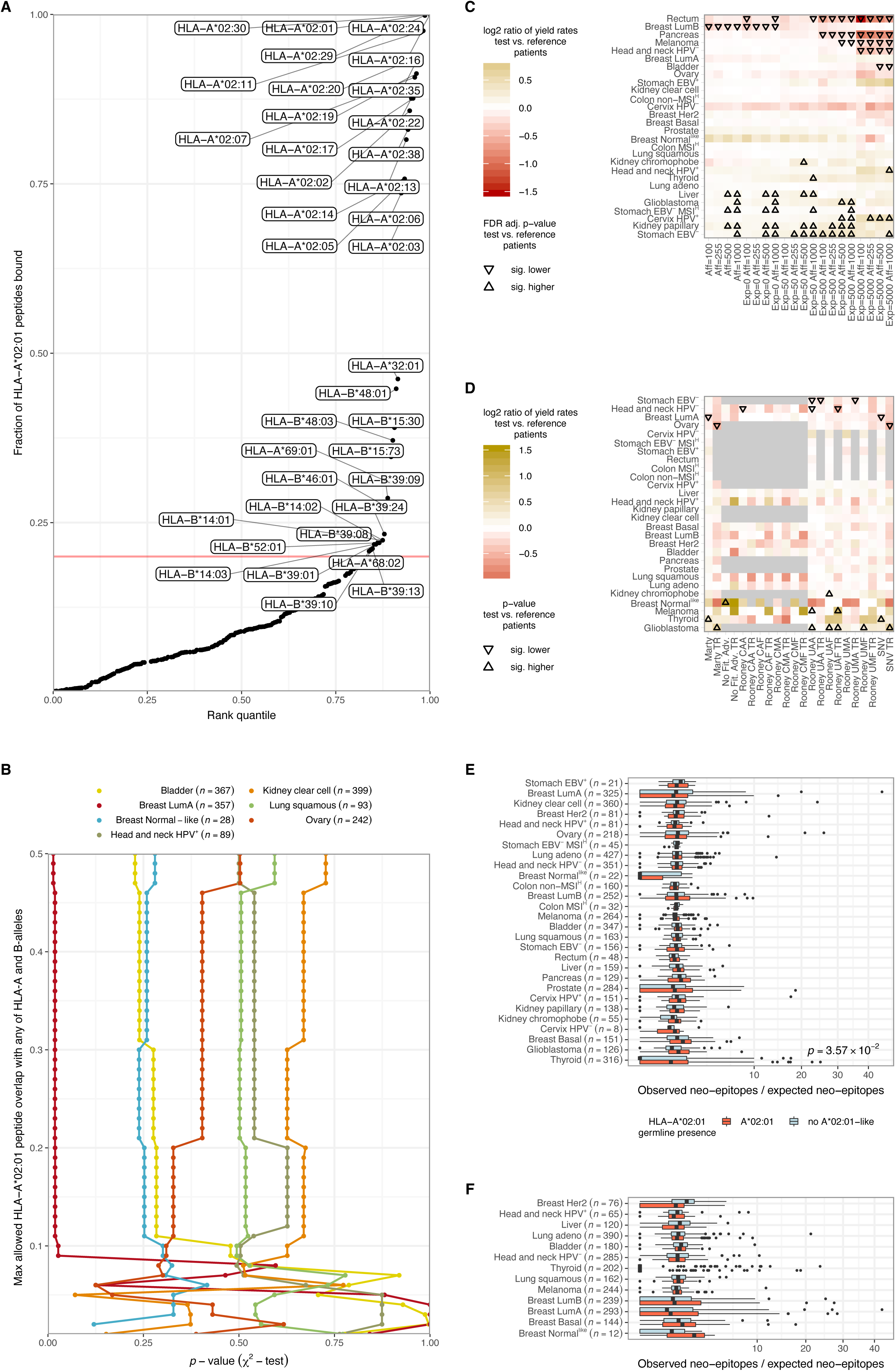
related to Figure 1: Extended analysis of genomic neoantigen depletion. **A.** Fraction of predicted HLA-A*02:01 neoantigens that is predicted to also be presented by any of the other detected HLA class I A and B alleles. Unless indicated otherwise (see B), HLA alleles predicted to present more than 20% of predicted HLA-A*02:01-neoantigens (as indicated by the red line) were labelled as ‘HLA-A*02:01-like’ in subsequent analyses, and samples expressing any of these alleles were excluded from the HLA-A*02:01-like negative reference group. **B.** The relationship between ‘the HLA-A*02:01-similarity threshold for patient inclusion in the HLA-A*02:01-like-negative reference set and the observed *p*-values in a meta-patient analysis of yield rate differences between the test samples and reference set samples is depicted per tumor type. To avoid over-plotting, seven randomly selected tumor types are shown. Note that an increase in stringency does not result in a systematic increase in statistical significance when comparing test and reference groups. **C.** Effect of different neoantigen prediction pipeline parameterizations on the detection of neoantigen depletion. Yield-rate differences in meta-patient analyses (see Figure 3A) are depicted. Triangles reflect comparisons for which FDR-adjusted *p*-values from two-sided Wilcoxon rank-sum tests were below 0.05. Upward and downward pointing triangles would point to genomic neoantigen depletion and to negative genomic neoantigen depletion, respectively. **D.** Effect of different modifications of neoantigen depletion analysis strategies on the detection of neoantigen depletion, while keeping neoantigen prediction parameters at the default settings. In the ‘Marty’-analyses, analysis is restricted to recurring oncogenic mutations as identified by Marty et al. (2017). Also shown are analyses in which mutations that can confer a fitness advantage are excluded (‘No Fit Adv.’), as in Figure 3D. In addition, 8 versions of analyses inspired by Rooney et al. (2015) are included. In this analysis, all permutations of three core settings were tested, denoted by three letters in the following format: ‘(C|U)(M|A)(F|H)’. Position 1 of the analysis identifier denotes the exclusion of variants based on clonality: (U)naware (no exclusion of subclonal mutations) vs. (C)lonal. Position 2 denotes the computation of conversion factors: (M)utational context specific vs. (A)specific. Position 3 denotes the neoantigen prediction pipeline filtering parameter: (F)ull 4-filter pipeline vs. HLA affinity filter (H) only. Finally, the SNV-restricted analysis also presented in Figure 3A is shown. Of all analyses, a T cell-resistance (TR) version is additionally shown, in which tumor samples that have at least one non-synonymous mutation in one of the 515 genes labelled as providing resistance to T-cell pressure^1^ are excluded. Unadjusted *p*-values from two-sided Wilcoxon rank-sum tests are shown by triangles in case of significance (*p* < 0.05). Upward and downward pointing triangles would point to genomic neoantigen depletion and to negative genomic neoantigen depletion, respectively. Missing analyses due to missing input data are indicated by a gray fill. **E.** Employing a mutation loss detection strategy inspired by Rooney et al. (2015), in which the expected neoantigen load is computed based on the synonymous mutation load and contrasted against the observed (predicted) neoantigen load. **F.** As in **E**, but also restricting the analysis to clonal mutations.

**Figure S4:**
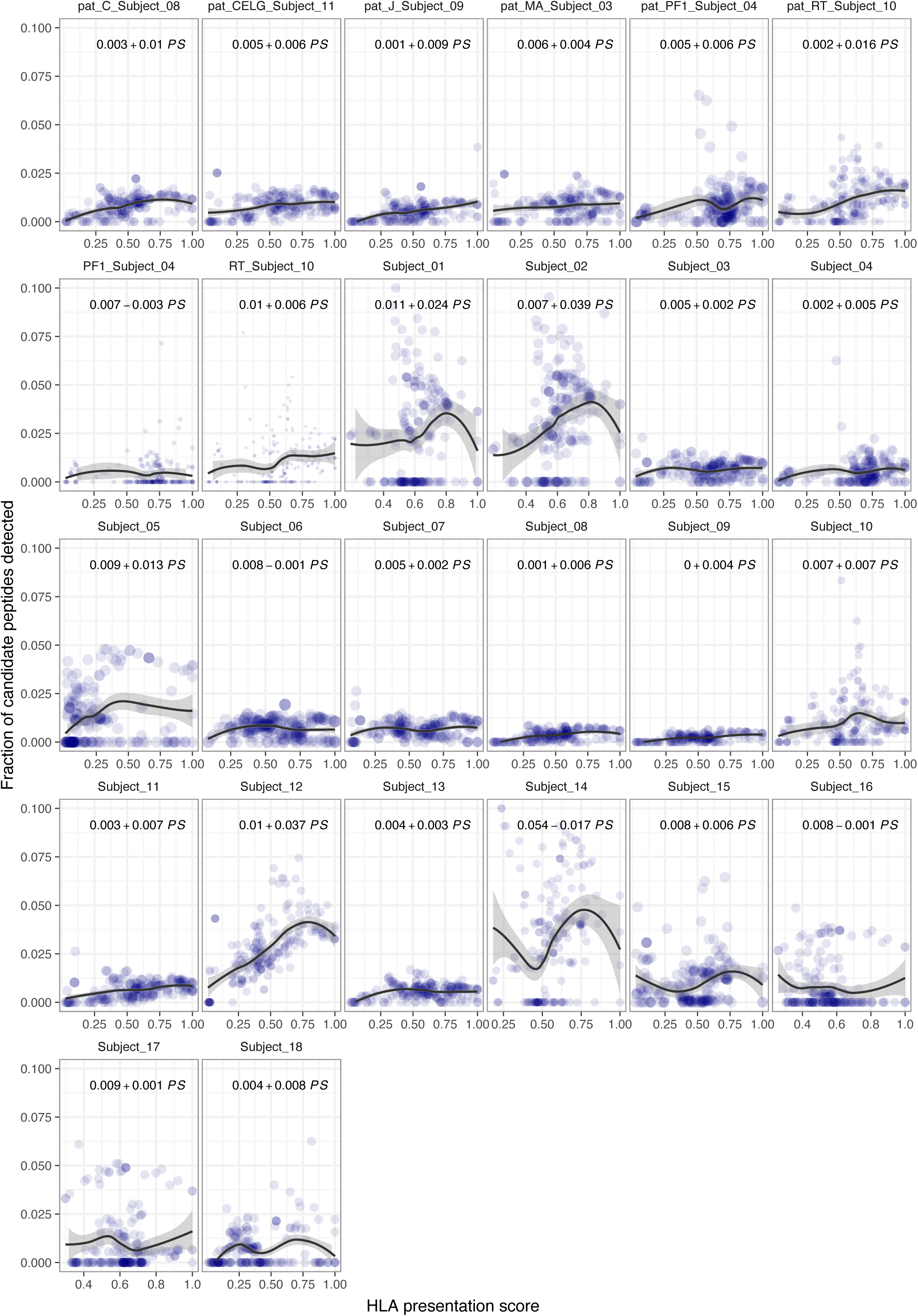
Mass-spec based validation of the presentation score for HLA-presentation capability. Each panel represents a single sample from the Pearson et al. (2016) study^2^ that assessed HLA-associated peptides by mass spectrometry. Each dot represents a focus allele. Horizontal axis: presentation score for this particular sample for a given focus allele, vertical axis: the fraction of expected peptides under 100% sensitive mass spec technology (i.e., with each peptide from every available protein detectable) that is detected. In general, HLA alleles that show a higher overlap with the predicted binding capacity of the HLA alleles that are expressed by a sample (i.e., a high presentation score) show higher fractions of detected peptides.

**Figure S5:**
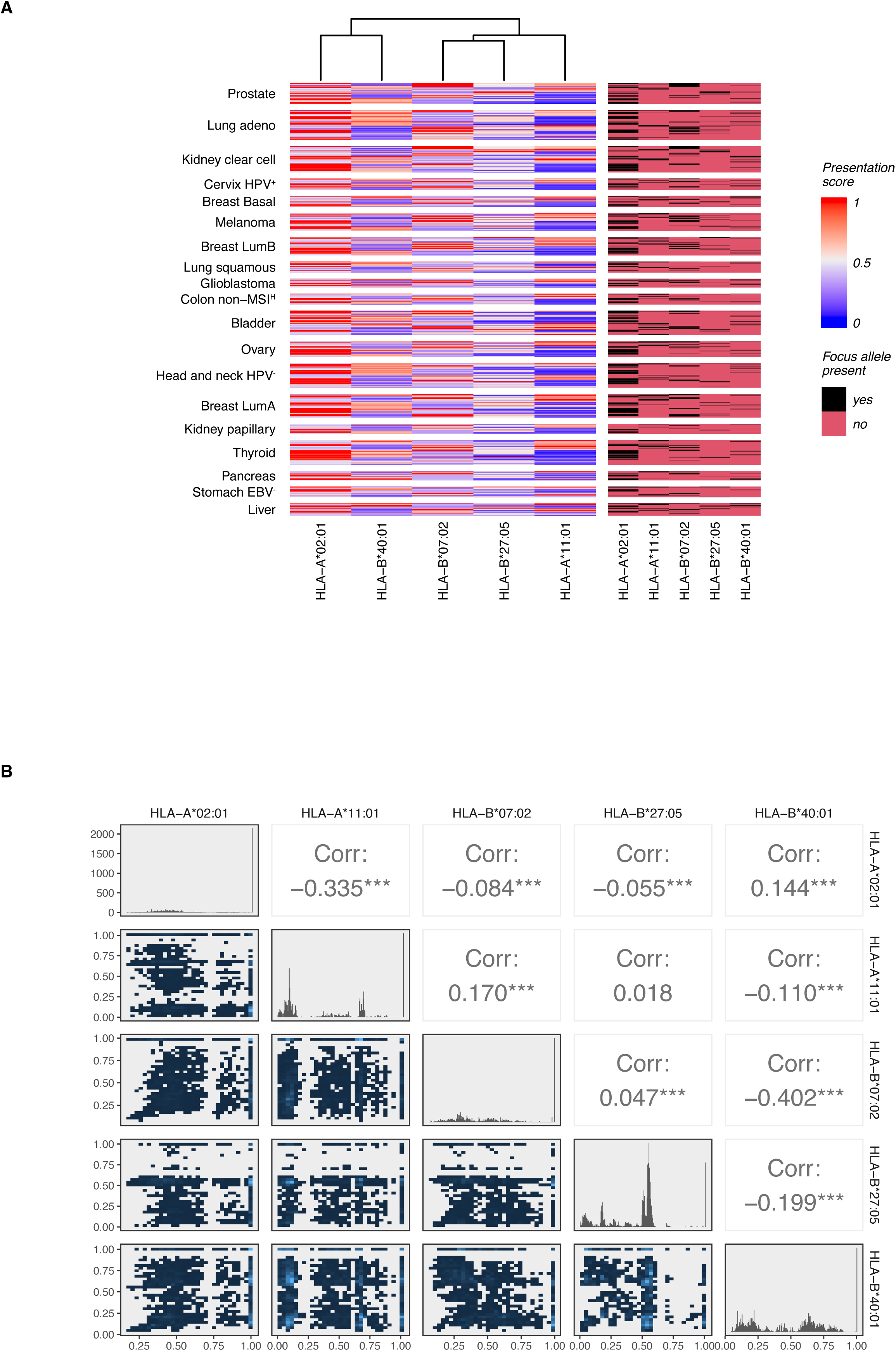
Presentation scores are highly variable across focus alleles. **A**. Heatmap of *h* across all included samples, computed across all 6 HLA class I alleles in a samples, not leaving out HLA alleles that are genomically lost. Left panel: raw presentation scores. Right panel: binary matrix indicating whether the focus allele is present in the set of a patient’s six HLA class I alleles. **B.** Pairwise correlation matrices of presentation scores. Diagonal elements (histograms) reflect univariate distributions. Sub-diagonal and supra-diagonal elements show a near-absence of correlation between focus alleles.

**Figure S6:**
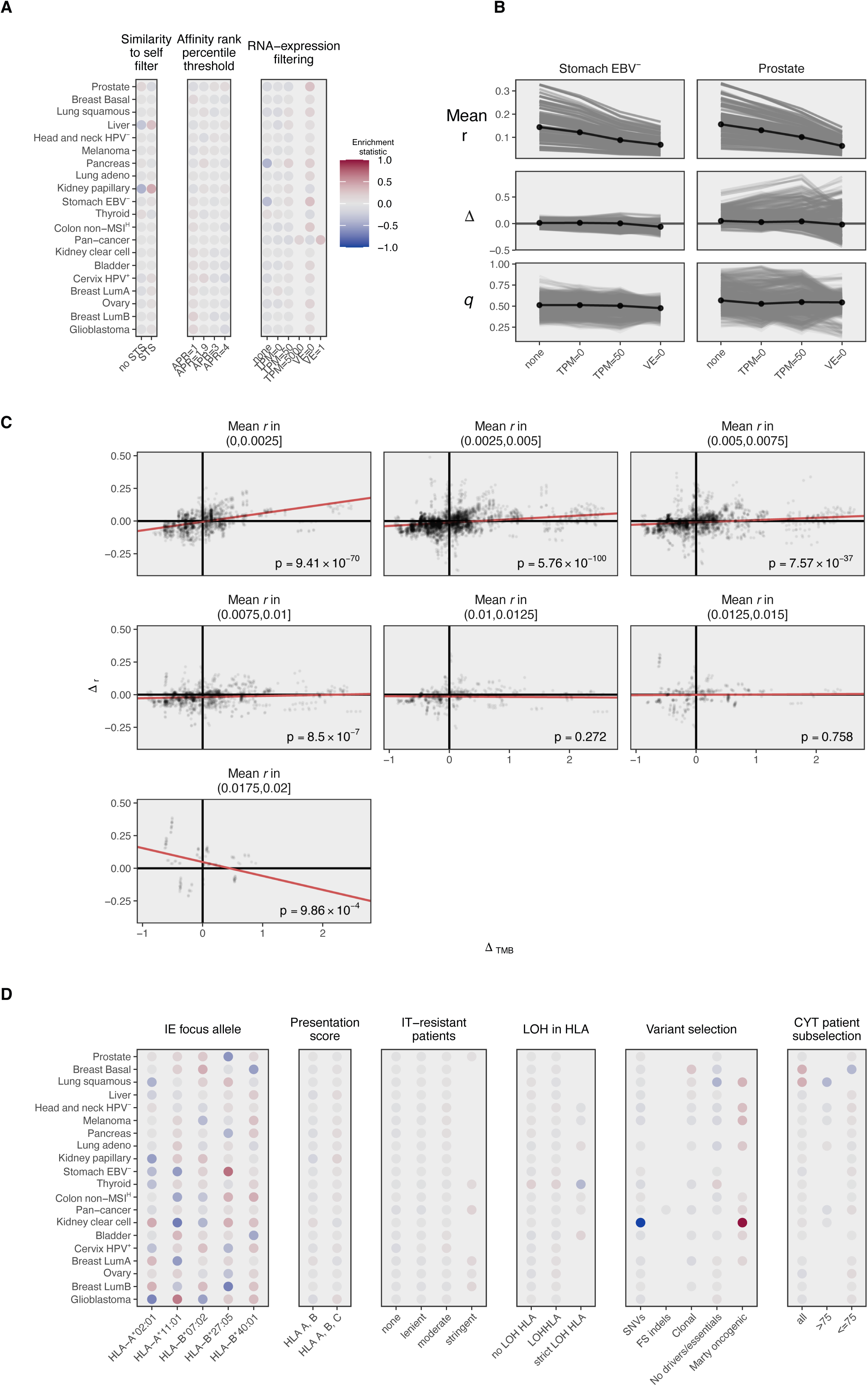
Efficient running of LOHHLA for large sample numbers. **A**. Relationship between the disk size of a TCGA BAM file and its read count content. The strong relationship here allows one to estimate the read content of BAM files from their file size, which, unlike the read count content, can be queried from the TCGA GDC API. Total read count is a required input variable for LOHHLA to run. **B.** Illustration of the robustness of HLA allele copy number estimates with regards to the LOHHLA minCoverage threshold that defines the minimal number of overlapping reads for a single genomic position to be considered robust enough for use in HLA allele copy number estimates. Each line corresponds to a single copy number estimate. **C.** Quantification of the robustness in B, using the coefficient of variation (CoV), visualized with violin plots. Most alleles have near-zero estimates, indicating robustness. **D.** The fraction of HLA allele copy number estimates (6 per patient) with CoV-estimates below the threshold

**Figure S7.**
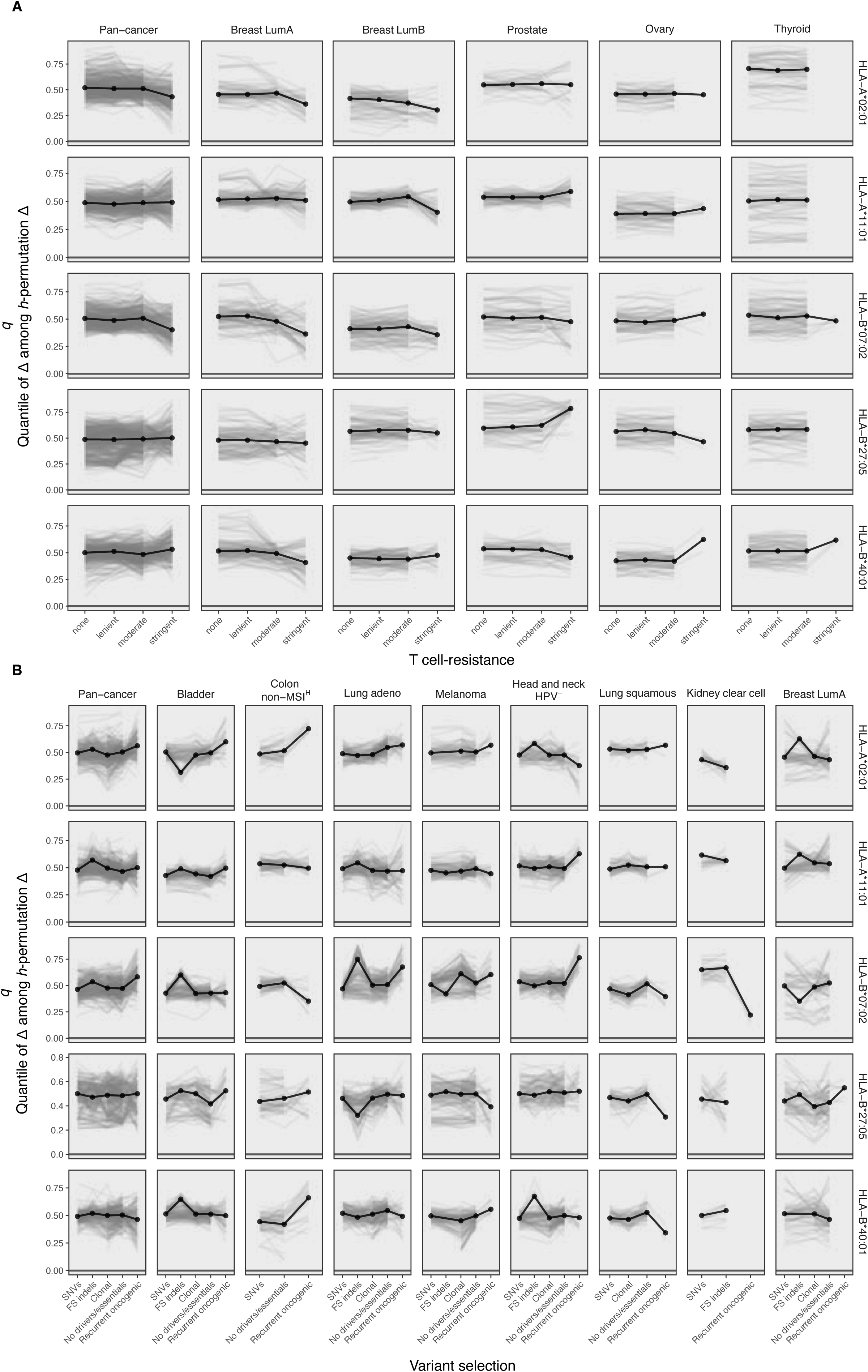
related to Figure 4: Sub-analysis of settings associated with Δ. **A**. As in Figure 4A, but after sorting of sub-analyses on Δ rather than *q*. Sub-analyses with the RNA-expression filter set to ‘VE=0’ (i.e., requiring > 0 / 10^6^ total reads to overlap the variant in order to be called expressed) appear enriched among the sub-analyses with the most negative Δ (consistent with neoantigen depletion). **B.** As in Figure 4D, but varying the RNA expression filtering setting on the horizontal axis. Variant-level filtering (VE=0) somewhat strongly affects the mean neoantigen yield rate (*r*) and also affects Δ. However, it does not affect *q* at all, indicating the effect on Δ is likely to be artefactual, see supplementary note 1. **C.** Evidence that stringent neoantigen filtering, resulting in low mean *r* across all patients, in combination with inhomogeneous distribution of *h*, causes Δ to be biased. Please see Supplemental Note 1 for associated reasoning. P-values in the lower right corners reflect the linear regression slopes of the Δ*_TMB_* vs. Δ*_r_* regression line. **D.** As in Figure 4B. but for settings associated with the manner in which the neoantigen depletion analysis is performed. Color scale as in A.

**Figure S8.**
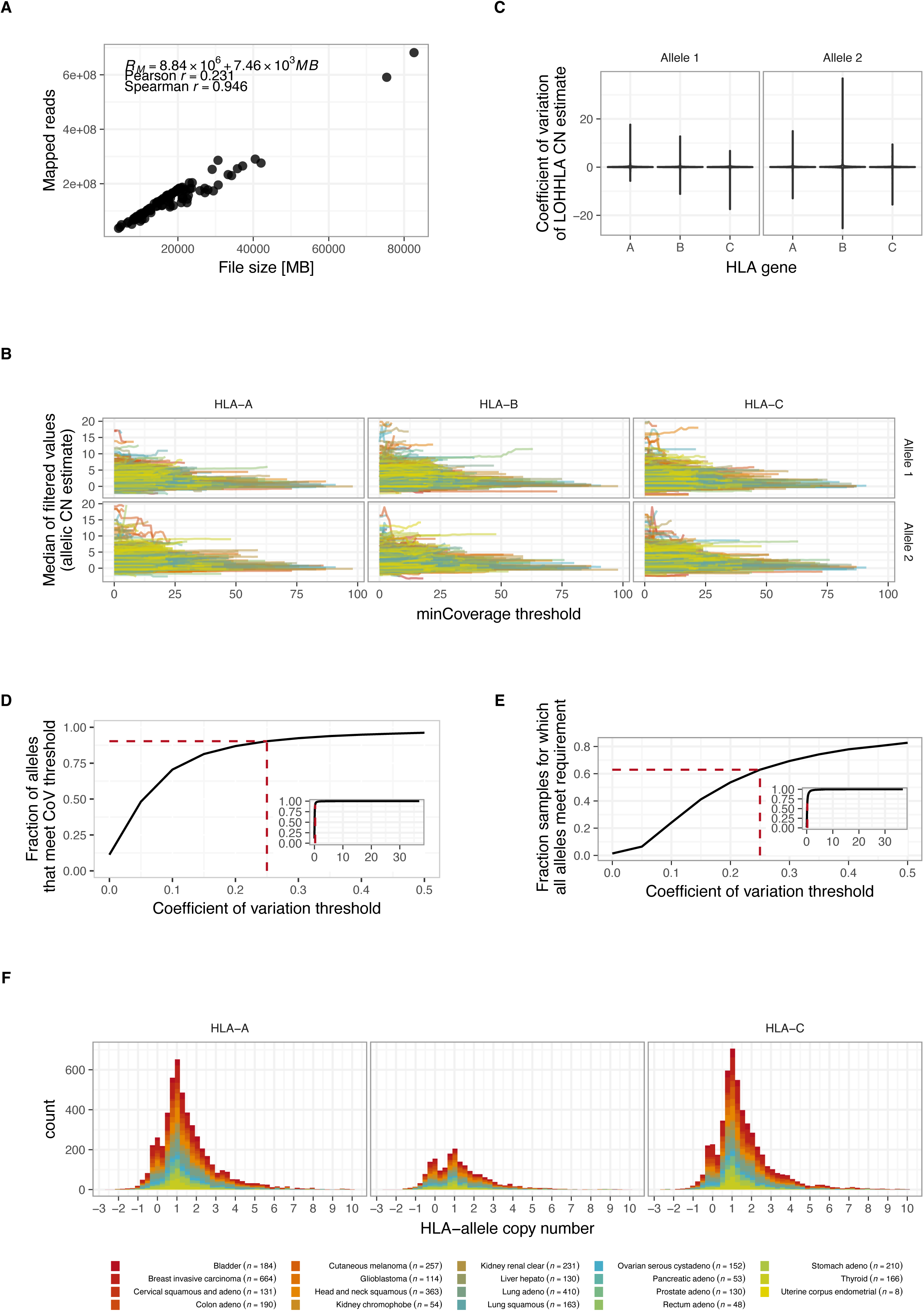
related to Figure 4: Trends of neoantigen depletion are not reproducible across focus alleles. **A.** As in Figure 4E, but split by focus allele and for all tumor types for which stringent filtering of T cell-resistant tumor samples was possible. **B.** As in Figure 4F, but split by focus allele and for all tumor types for which the recurrent oncogenic variant selection was possible.

