## Supplementary Note 1 for "Lack of detectable neoantigen depletion in treatment-naïve cancers"

### Supplemental Note 1

Testing for neoantigen depletion settings that were associated with $\Delta$ (rather than $q$, as used in the main text), we found that stringent RNA expression filtering on the variant level on average led to a negative $\Delta$ for a majority of tumor types (Figure S7A). Selecting and plotting groupings of identically parameterized sub-analyses, save for the RNA expression filtering setting, confirmed the observation of an effect on Δ and not $q$ (Figure S7B).

Interpreting this result, we imagined that stringent neoantigen filtering might enrich for negative Δ but not low *q* through lowering the mean neoantigen yield rate ($r$). As low-TMB patients will be more likely to be predicted to have 0 neoantigens, appearing fully depleted of neoantigens, severe reduction of $r$ could render the overall analysis sensitive to imbalances in the distribution of patients over the presentation score ($h$*)*. This is because any range of $h$ that is relatively sparse in patients will more likely appear to be fully depleted of neoantigens. When relatively few patients populate the lower range of $h$ ($h\in[0,\sim.25]$), Δ could then be biased upwards (i.e., indicating nonsensical enrichment of neoantigens with enhanced neoantigen presentation). Similarly, Δ will be biased downwards with relative patient sparsity at the upper range of $h$ ($h\in[\sim.75, 1]$).

To test this hypothesis, we first computed the mean relative difference in TMB between $h=1$ and $h=0$ patients ($\Delta_{TMB}$), much like we did in our primary regression of $h$ against $r$ (Δ), to directly quantify the degree with which $h$ and TMB are correlated. This demonstrated that $\Delta_{TMB}$ is strongly associated with Δ (Figure S7B), especially for sub-analyses for which the mean $r$ is low, indicating that imbalance in the distribution of $h$ at least partially explains $\Delta$ (Figure S7C). As $h$-permutation does not modify the overall $h$-distribution, permutation $\Delta$s will retain whatever degree of bias that is already present in the original data, such that $q$ is the more robust and informative of the two statistics to assess.

We conclude that, through lowering $r$, in combination with inhomogeneous distribution of $h$, variant level expression filtering likely lowered $\Delta$ but not $q$ in a manner that is independent from biology.
